# Loss of Sister Kinetochore Co-orientation and Peri-centromeric Cohesin Protection after Meiosis I Depends on Cleavage of Centromeric REC8

**DOI:** 10.1101/2020.02.06.935171

**Authors:** Sugako Ogushi, Ahmed Rattani, Jonathan Godwin, Jean Metson, Lothar Schermelleh, Kim Nasmyth

**Author notes:** Correspondence (K.N.).

## Abstract

Protection of peri-centromeric REC8 cohesin from separase and sister kinetochore attachment to microtubules emanating from the same spindle pole (co-orientation) ensure that sister chromatids remain associated after meiosis I. Both features are lost during meiosis II, when sister kinetochores bi-orient and lose peri-centromeric REC8 protection, resulting in sister chromatid disjunction and the production of haploid gametes. By transferring spindle-chromosome complexes (SCCs) between meiosis I and II cells, we have discovered that both sister kinetochore co-orientation and peri-centromeric cohesin protection depend on the SCC and not the cytoplasm. Moreover, the catalytic activity of separase at meiosis I is necessary not only for converting kinetochores from a co-to a bi-oriented state but also for deprotection of peri-centromeric cohesin and that cleavage of REC8 may be the key event. Crucially, we show that selective cleavage of REC8 in the vicinity of kinetochores is sufficient to destroy co-orientation in univalent chromosomes, albeit not in bivalents where resolution of chiasmata through cleavage of Rec8 along chromosome arms may also be required.

## Introduction

The production during meiosis of haploid gametes from diploid germ cells is only possible because two rounds of chromosome segregation occur without an intervening round of DNA replication. As during mitosis, meiotic DNA replication is accompanied by establishment of cohesion between sister DNAs, which involves entrapment of DNAs inside cohesin rings. This ensures that just a single reciprocal recombination event between homologous non-sister DNAs, creating chiasmata, joins all four homologous chromatids together, forming bivalent chromosomes (Nasmyth, 2015; Watanabe, 2012).

During meiosis I, sister kinetochores act as a single unit (co-orientation) and as a consequence, pairs of maternal and paternal kinetochores, but not sister kinetochores, are pulled in opposite directions by microtubules. Under these circumstances the tension required to stabilize kinetochore-microtubule interactions depends solely on sister chromatid cohesion distal to chiasmata. The first meiotic division is triggered by cleavage of cohesin’s REC8 subunit along chromosome arms by a thiol protease called separase, which resolves chiasmata and creates dyads (Kudo et al., 2006). The two chromatids of these dyads are held together through cohesion between their peri-centromeres and sister kinetochores are pulled in opposite directions (bi-orientation) until separase cleaves the remaining cohesin molecules during the second round of chromosome segregation. (Kitajima et al., 2006; Riedel et al., 2006; Tang et al., 2006). Two key features therefore distinguish the first meiotic division from the second and from mitosis, namely co-orientation of sister kinetochores and protection of peri-centromeric cohesin. How these are conferred and how they are both subsequently lost, so that during meiosis II microtubules pull sister kinetochores in opposite directions (bi-orientation) and separase cleaves peri-centromeric cohesin thereby converting dyads into chromatids, is poorly understood.

Peri-centromeric cohesin is protected from separase during meiosis I by proteins belonging to the Shugoshin/Mei-S332 family which form a highly conserved homodimeric parallel coiled-coil that binds protein phosphatase 2A, PP2A-B56 (Xu et al., 2009). Protection arises because cleavage by separase of cohesin’s REC8 kleisin subunit requires REC8’s prior phosphorylation, which is removed by PP2A (Attner et al., 2013; Brar et al., 2006; Katis et al., 2010). Recruitment of PP2A by shugoshin is necessary for peri-centromeric cohesion protection in mammals (Lee et al., 2008; Rattani et al., 2013) as well as in budding and fission yeasts (Kitajima et al., 2006; Riedel et al., 2006), suggesting that this mechanism is conserved throughout eukaryotes. What is less clear is how shugoshin ensures that PP2A de-phosphorylation of REC8 selectively within peri-centromeric chromatin and how this ability is lost after the first meiotic division.

Equally mysterious is the mechanism by which protection of peri-centromeric cohesin from separase is lost by the time cells initiate anaphase II. In budding yeast, degradation by APC/C^Cdc20^ of its sole shugoshin ortholog Sgo1 at the onset of anaphase II, a phenomenon that does not occur at the equivalent stage of meiosis I, ensures that Sgo1 and PP2A are removed by the time separase is activated (Jonak et al., 2017). However, SGOL2, which confers protection during meiosis I in mammals, does not possess this regulatory feature (Marston, 2015). Moreover, ectopic accumulation of Sgo1 in the vicinity of centromeres in fission yeast is insufficient to protect peri-centromeric REC8 during meiosis II (Gregan et al., 2008). Likewise, the suggestion that protection is destroyed by tension created within peri-centromeric chromatin through bi-orientation of sister kinetochores during meiosis II fails to explain adequately how such tension would actually ablate shugoshin’s activity (Lee et al., 2008). It also fails to explain why bi-orientation of sister kinetochores in yeast monopolin mutants is not accompanied by de-protection (Yokobayashi & Watanabe, 2005). Factors such as I2PP2A and cyclin A have been postulated to have a role in de-protection in mammals (Touati et al., 2012; Wassmann, 2013) but it remains unclear how they might function.

Unlike the protection of centromeric cohesin, the mechanism responsible for sister kinetochore co-orientation during meiosis I appears to differ among species. (McKinley and Cheeseman, 2016). In budding yeast whose point centromeres contain a single CENPA nucleosome, co-orientation during meiosis I depends on a monopolin complex, the core of which is the V shaped Csm1 heterodimer, that thought to confer co-orientation by cross-linking sister kinetochores (Corbett and Harrison, 2012; Rabitsch et al., 2003; Sarkar et al., 2013). Because Csm1 has little or no role in co-orientation in the fission yeast *S.pombe* (Gregan et al., 2008) and indeed is absent from metazoan genomes (Plowman et al., 2019), some other mechanism must confer co-orientation in eukaryotes with regional centromeres. In fission yeast, a meiosis-specific version of cohesin containing a Rec8 kleisin subunit is necessary to prevent bi-orientation during meiosis I (Watanabe and Nurse, 1999; Sakuno et al, 2009). This suggests that cohesin mediates co-orientation by holding sister centromeres together. Though plausible, this hypothesis has never been rigorously proven. In an attempt to target specifically the centromeric cohesin population for cleavage using fission yeast strains expressing a CenpC-TEV fusion and Rec8 containing TEV sites, there was little or no evidence that centromeric cohesin was more depleted than peri-centromeric cohesin (Yokobayashi & Watanabe, 2005).

Another important meiotic regulatory factor both in fungi and metazoa is a family of proteins related to Spo13 in budding yeast (Wang et al., 1987). These include Moa1 in fission yeast (Yokobayashi & Watanabe, 2005) and Meikin (Kim et al., 2015) in mammals. Though not highly conserved in amino-acid sequences, all members of the family are expressed exclusively during meiosis and have the property of recruiting Polo-like kinases to kinetochores. Because their ablation compromises protection of peri-centromeric cohesion by Shugoshin/Mei-S332 proteins (Katis et al., 2004a; Klapholz & Esposito, 1980; Lee et al., 2004; Shonn et al., 2002) and regulation of the anaphase-promoting complex/cyclosome (APC/C) activity (Katis et al., 2004b) as well as co-orientation (Galander et al., 2019; Kim et al., 2015; Matos et al., 2008; Miyazaki et al., 2017), these Spo13-like proteins should be viewed as factors that regulate directly or indirectly numerous properties of meiotic centromeres.

To address the mechanisms conferring co-orientation and peri-centric protection in mouse oocytes, we have adopted a technique developed more than 40 years ago with grasshopper spermatocytes, namely the transfer of chromosomes and their spindles from meiosis I to meiosis II cells and vice versa (Nicklas, 1977; Paliulis and Nicklas, 2000). Our findings confirm that both co-orientation of sister kinetochores and peri-centromeric cohesin protection are conferred by the state of chromosomes and their spindles and not by the nature of meiosis I cytoplasm. By combining cytological and genetic manipulations, we show that separase cleavage activity is required to convert chromosomes from a meiosis I to a meiosis II state, and that conversion is most likely mediated by cleavage of a specific pool of REC8-containing cohesin associated with centromeric DNA. Our findings suggest that centromere-specific cohesin not only confers the co-orientation of sister kinetochores but more unexpectedly also helps shugoshin (SGOL2) protect from separase peri-centromeric cohesin located many megabases away (Vissel and Choo, 1989). Centromere-specific cohesin is also protected by SGOL2 upon separase activation during meiosis I. However, unlike peri-centromeric cohesin, which survives until cells embark on the second meiotic division, the protection of centromeric cohesin by shugoshin only lasts until late telophase I, whereupon REC8 cleavage induces sister kinetochores to split into their component parts, setting the scene for their subsequent bi-orientation during meiosis II.

## Results

### Dyads retain their MII character when transferred to MI oocytes

To address whether factors associated with the spindle-chromosome complex (SCC) or those within the cytoplasm confer co-orientation and/or peri-centromeric cohesion protection, we used microsurgery to transfer SCCs between oocytes. We first addressed how dyads isolated from metaphase II oocytes (the donor cells) behave when placed inside meiosis I oocytes (the host cells). SCCs from metaphase II oocytes, which are surrounded by membrane, were placed in contact with metaphase I oocytes and fused with them using envelopes from Hemagglutinating Virus of Japan (HVJ-E) (Fig. 1A). The overall fusion success rate was 94% (866/920). To observe the behaviour of chromosomes from the donor and host cell, both were injected at the germinal vesicle (GV) stage with mRNAs encoding histone *H2b* fused to *mCherry* (*H2b-mCherry*), which labels entire chromosomes, and *Cenp-C* fused to *eGfp* (*eGfp-CenpC*), which labels kinetochores. To assess the effect of small amounts of cytoplasm transferred along with MII SCCs, we also fused MI oocytes with cytoplasts from MII oocytes that had a similar volume to SCCs (MIIcyt+MI). Importantly, neither the frequency nor timing of chromosome segregation differed substantially between intact unmanipulated oocytes (86%, 14/16, 534±79 min), MIIcyt+MI oocytes (76%, 13/17, 536±61 min), or MII-SCC+MI oocytes (86%, 37/43, 563±94 min). The fusion procedure therefore had little or no adverse effect on meiotic progression (Figs. 1B, and S1A).

**Figure 1.**
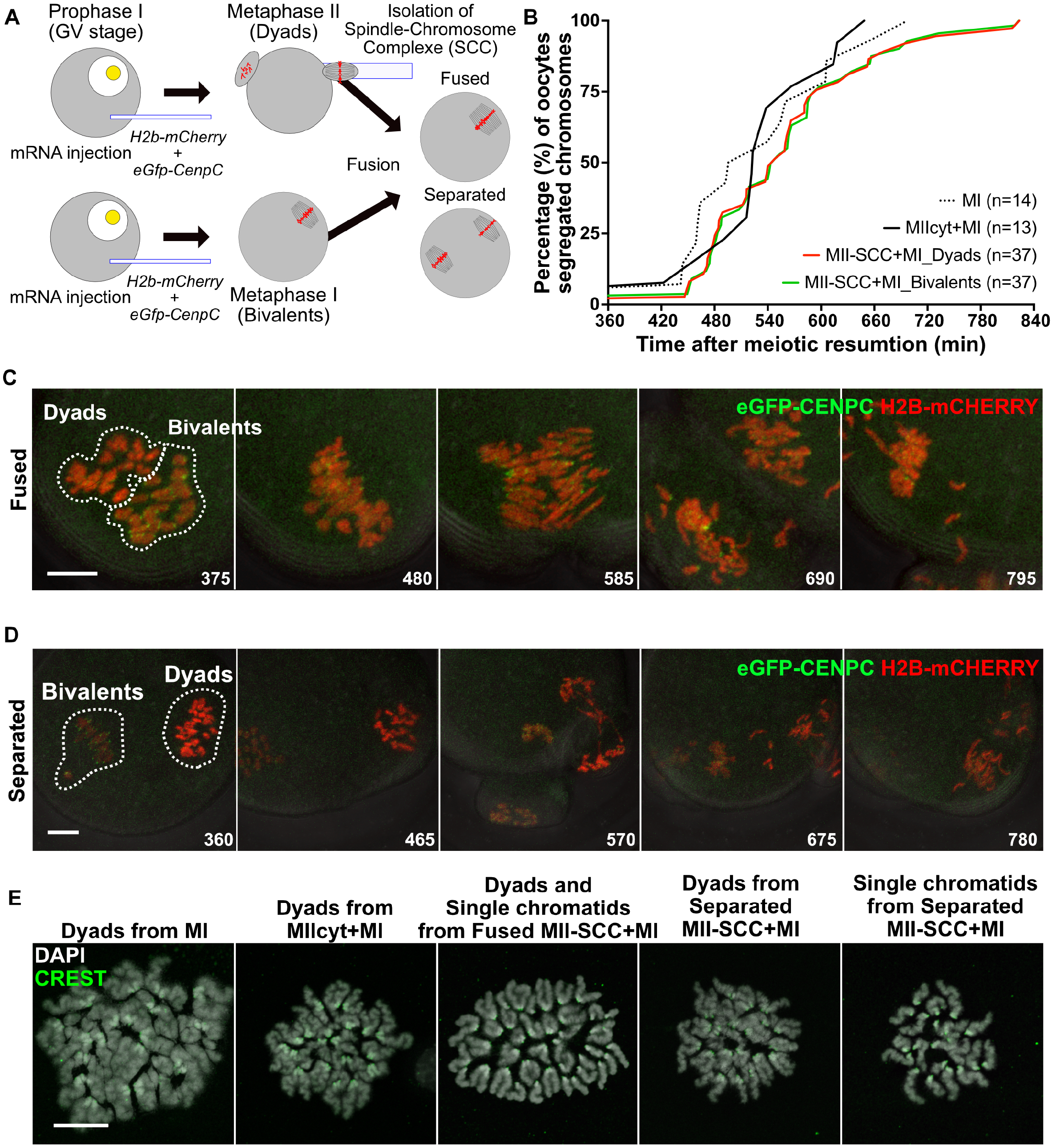
Either Sister Kinetochore Bi-orientation or De-protection of Cohesion Is Conferred by the Nature of Dyads Itself rather than the Surrounding Factors Coming from Cytoplasm. (A) Schematic of an experiment showing that a spindle chromosome complex (SCC) containing dyads from an oocyte at the second metaphase (MII) is fused with an oocyte at the first metaphase (MI), and a resulting oocyte is called MII-SCC+MI. (B) Segregation timing of chromosomes in MII-SCC+MI. As a technical control, an MII cytoplasm was fused with an MI oocyte (MIIcyt+MI). n, the numbers of oocytes measured in more than three independent experiments. (C) Representative stills from live cell imaging of chromosome segregation in MII-SCC+MI, whose MII-SCC was fused with a host MI spindle. Chromosomes were labelled by H2B-mCHERRY (red) and kinetochores were visualized by eGFP-CENPC (green). Numbers indicate the time after meiotic resumption (min). Bar, 10 µm. (D) Representative stills from live cell imaging of chromosome segregation in MII-SCC+MI without spindle fusion as in Figure 1C. Bar, 10 µm. (E) A representative image of chromosome spread showing single chromatid formation from transferred dyads after the first meiotic division. The antibody is shown in the panel (green). DNA was stained with DAPI (grey). Bar, 15 µm.

In 61% (26/43) of cases, MII-SCCs formed a single spindle together with the host MI-SCCs. In 88% of these (23/26), the dyads transferred to MI oocytes disjoined to form individual chromatids at the same time as bivalents were converted to dyads (Fig. 1C, E & Movie 1). Three oocytes failed to undergo chromosome segregation and arrested in metaphase I. In those oocytes in which the MII-SCC remained separate from the host MI-SCC (39%, 17/43), the dyads disjoined, to form individual chromatids and this process coincided with conversion of bivalents to dyads on the host MI spindle (Fig. 1D, E & Movie 2). This simultaneous double chromosome segregation took place in 82% (14/17) of these oocytes. Most of these double divisions were accompanied by cytokinesis, producing two polar bodies. However, in 2 out of 17 (12%) only one of the two SCCs triggered cytokinesis.

Two conclusions can be drawn from these findings. First, despite the presence of an MI cytoplasm conducive to the co-orientation of maternal and paternal kinetochores associated with bivalent chromosomes, dyad kinetochores bi-orient just as they do in MII oocytes. An MI cytoplasm cannot therefore induce the co-orientation of sister kinetochores associated with dyads. Second, neither the cytoplasm of MI oocytes nor the adjacency of MI bivalents can protect from separase the peri-centromeric cohesin holding dyads together. Both the propensity to bi-orient and the susceptibility of peri-centromeric cohesin to separase are therefore properties associated with SCCs and not conferred by cytoplasmic state.

One potential reason why bi-orientation persists after transferring dyads to MI oocytes is that the connections between kinetochores and microtubules established in MII persist throughout the transfer process and are subsequently maintained until anaphase is initiated. To exclude this possibility, we transferred MII SCCs into oocytes at the germinal vesicle (GV) stage (MII-SCC+GV), a procedure that leads to dissolution of pre-existing spindles or microtubule-kinetochore attachments. The fused oocytes were cultured for a further 2-4 hours before they were released from their GV arrest and allowed to undergo meiosis I (Fig. S1B, C, Movies S1 & S2). Under these conditions, the transferred dyads and the host bivalents invariably (20/20) aligned on a single spindle but anaphase was initiated in only 25% of oocytes (5/20) compared to 55% (6/11) when MII cytoplasts were fused to GV oocytes (MII-SCC+GV). The failure of many oocytes to undergo anaphase was usually accompanied by the presence of rare dyads that failed to associate with the spindle, which presumably led to SAC activation. Importantly, such dyads were never observed in oocytes that did undergo anaphase. Furthermore, even in oocytes that failed to undergo anaphase, those dyads that had associated with spindles (which was the vast majority) clearly bi-oriented (Fig. S1D & Movie S3). Crucially, in the five oocytes that actually underwent anaphase, their dyads disjoined to form individual chromatids (Fig. S1C), confirming that their bi-orientation was functional. Thus, the kinetochores of dyads fail to co-orient even though all kinetochore attachments are made anew when GV oocytes enter meiosis I.

### Bivalents retain their character when transferred to MII oocytes

We next addressed whether the kinetochore co-orientation and peri-centromeric cohesin protection associated with bivalent chromosomes associated with MI SCCs is retained when they are transferred to MII oocytes (MI-SCC+MII) that are then triggered to undergo meiosis II division by adding strontium (Fig. 2A). Because the movements associated with cytokinesis hinder chromosome imaging under these conditions, cytokinesis was prevented by adding cytochalasin B at 5 µg/ml. To assess the effect merely of transferring a small amount of MI cytoplasm, we also prepared MII oocytes that had been fused with an MI cytoplast having a similar volume as an SCC from MI (MIcyt+MII). Chromosome segregation occurred in 24/39 (62%) MII oocytes fused with MI-SCC (MI-SCC+MII), a rate comparable with unfused MII oocytes (63%, 5/8) or oocytes fused to MI cytoplasts (MIcyt+MII) (75%, 3/4). There was a modest delay in MI-SCC+MII oocytes (Figs. 2B, S2A and B). Considering that oocyte activation is triggered by a rapid influx of intracellular Ca^2+^ concentration released from endoplasmic reticulum in oocyte cytoplasm (Miao et al., 2012; Wakai and Fissore, 2013), this delay could be caused by a lower ratio of cytoplasm to chromosomes in MI-SCC+MII oocytes. In 13/39 (33%) MI-SCC+MII oocytes, the introduced SCC inter-mingled closely with the host SCC, forming a single spindle. Under these circumstances, 12/13 (92%) underwent chromosome segregation, during which MI bivalents were converted to dyads while MII dyads were converted to individual chromatids (Fig. 2C, E & Movie 3). In the rest (26/39), the MI-SCC remained separate from the host MII-SCC. Of these, chromosome segregation took place in 46% (12/26), with bivalents being converted to dyads simultaneously with the conversion of dyads to individual chromatids (Fig. 2D & Movie 4). Thus, both kinetochore co-orientation and protection of peri-centromeric cohesion persist when bivalents are exposed to an MII cytoplasm.

**Figure 2.**
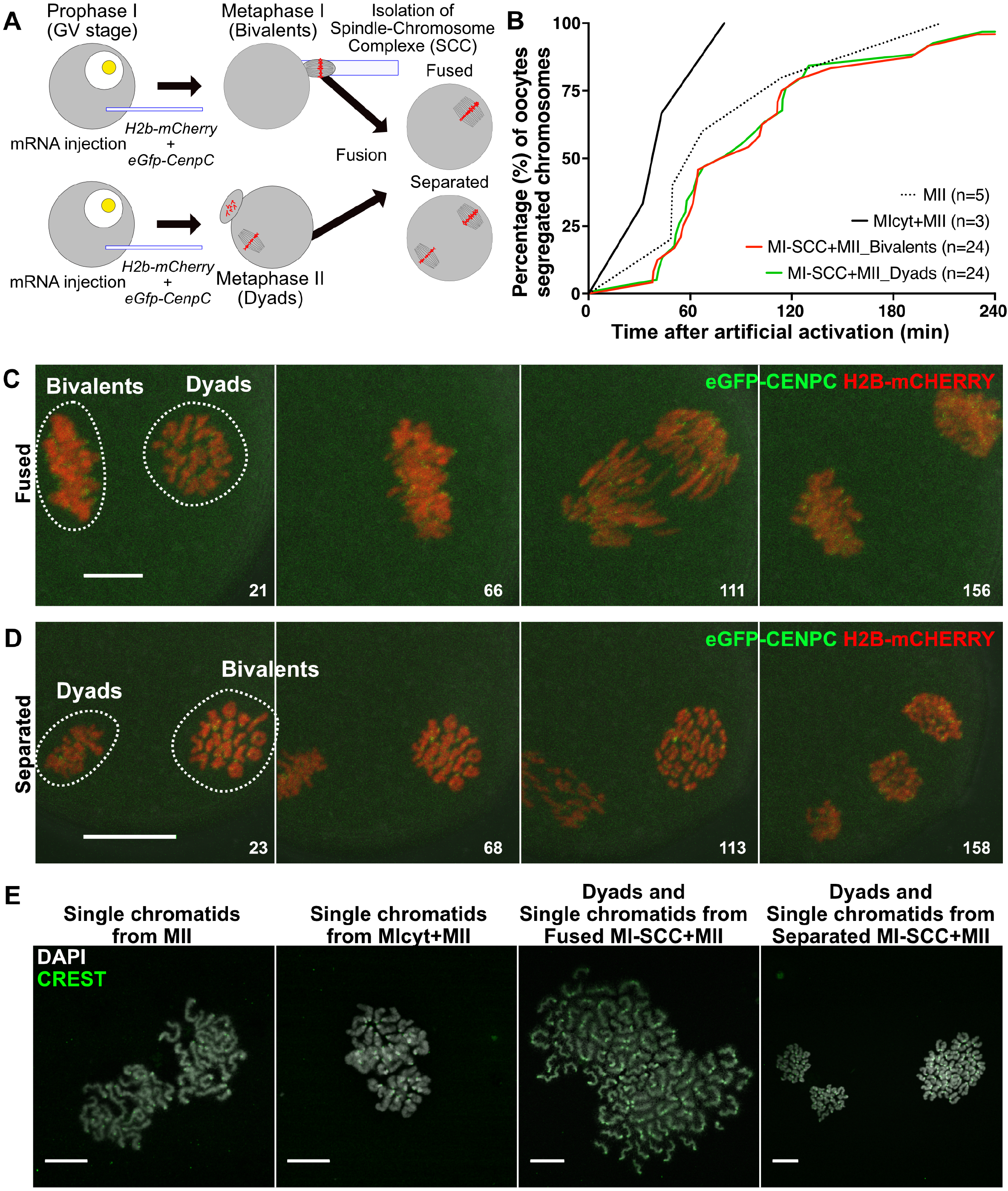
Determination of Protection of Cohesion or Sister Kinetochore Co-orientation Is Granted by Bivalents rather than the Factors from Cytoplasm. (A) Schematic of experiments showing an SCC containing bivalents is fused with an oocyte at MII (MI-SCC+MII). (B) Timing of bivalent segregation in MI-SCC+MII after artificial strontium activation. As a technical control, an MI cytoplasm was fused with an MII oocyte (MIcyt+MII) and artificially activated. n, the numbers of oocytes measured in more than two independent experiments. (C) Representative stills from live cell imaging of chromosome segregation in an MI-SCC+MII, whose MI-SCC was fused with a host MII spindle as in Figure 1C. Numbers indicate the time after artificial activation (min). Bar, 10 µm. (D) Representative stills from live cell imaging of chromosome segregation in MII-SCC+MI without spindle fusion as in Figure 2C. Bar, 20 µm. (E) A representative image of chromosome spread showing dyad formation from transferred bivalents at anaphase II (AII). The antibody is shown in the panel (green). DNA was stained with DAPI (grey). Bars, 15 µm.

### Separase cleavage at meiosis I is necessary for loss of peri-centromeric cohesin protection and kinetochore co-orientation

The previous experiments suggest that a change in the state of chromosomes rather than that of cytoplasm is responsible for the change in the behaviour of kinetochores and peri-centromeric cohesin upon completion of meiosis I. REC8 cleavage by separase is a well-known chromosomal process during MI-MII transition. To test whether this might play a role, we used an oocyte-specific knockout mouse line *Zp3-Cre Espl1* (*separase) (f/f)*, hereafter refer to as *Sep* (-/-), and a *separase* (f/f) mouse line as a control (*Sep* (+/+), Kudo et al., 2006). As previously reported, the lack of chiasmata resolution in oocytes from *Sep* (-/-) mice is accompanied by a failure to extrude the first polar body (in 43/44 or 98% of oocytes). Polar body extrusion was rescued in 57/105 (55%) oocytes by injection of separase mRNA, a success rate comparable with wild type controls (61%, 17/28). Injection of mRNAs encoding a catalytically inactive separase (C2028S) also rescued polar body extrusion in 124/224 oocytes (55%) but did not restore chiasmata resolution as previously shown (Fig S3A, Kudo et al., 2006). Thus, oocytes specifically defective in cleavage activity but no other separase functions, undergo most if not all cell cycle events that normally accompany APC/C activation during meiosis I, producing cells that contain bivalent chromosomes instead of dyads. To address whether separase cleavage activity is required to convert chromosomes from an MI state to an MII state, the SCC from a *Sep* (-/-) *C2028S* oocyte that had extruded a polar body at the first meiotic division was transferred to a wild-type MI oocyte, whose bivalents had been removed (Fig 3A). Crucially, the bivalents on this SCC were converted to dyads with kinetics similar to those of *Sep* (+/+) oocytes (Fig. 3B-D & Movies 5-7). Moreover, the dyads created were subsequently converted to individual chromatids when a second meiotic division was triggered through artificial activation (Fig S3B-D, Movies S4 & S5). This result implies that cleavage of an SCC-associated protein by separase is necessary to destroy co-orientation and protection of peri-centromeric cohesion.

**Figure 3.**
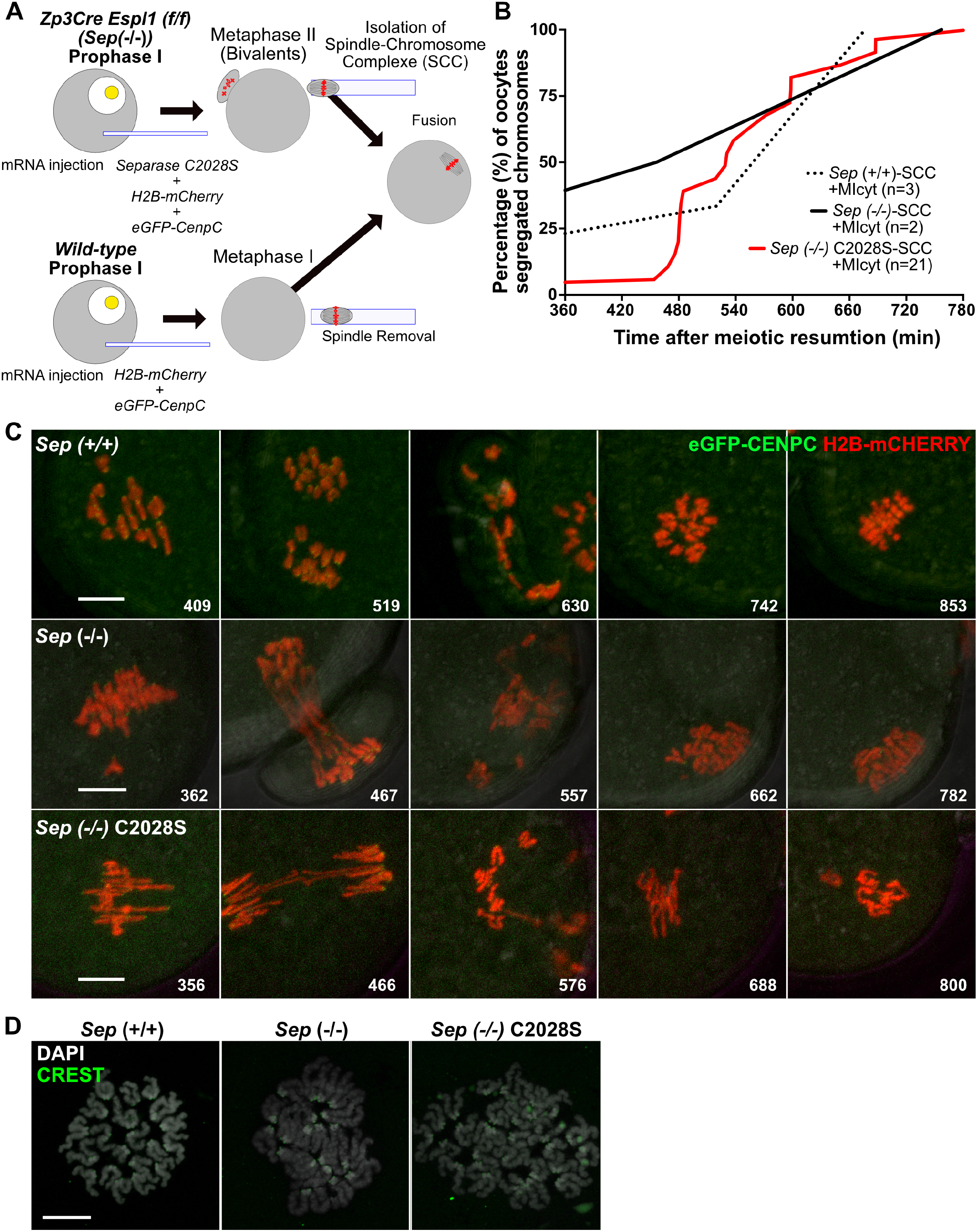
Cleavage of Cohesin by Separase Is Required for Changing the Status of Bivalents for Sister Kinetochore Orientation and for a Protection/De-protection of Peri-centromeric Cohesion. (A) Schematic of experiments showing that an SCC containing bivalents that have experienced all pathways during MI-MII transition, except REC8 cleavage by ESPL1 (separase), is fused with an MI cytoplast. (B) Timing of bivalent segregation in MI cytoplasts. We prepared three kinds of bivalents whose SCC were fused with MI cytoplasts (MIcyt). One is bivalents from *separase (f/f)* mouse line for control (*Sep (+/+)*-SCC+MIcyt), second is those from *Zp3Cre separase* (f/f) oocytes (*Sep (-/-)*-SCC+MIcyt) and the last is those from *Zp3Cre separase* (f/f) oocytes induced only cytokinesis by expression of catalytic-dead version of separase C2028S (*Sep (-/-)* C2028S-SCC+MIcyt). n, the numbers of oocytes measured in more than two independent experiments. (C) Representative stills from live cell imaging showing segregation of transferred bivalents in MI cytoplasts as in Figure 1C. *Sep (+/+)*: *Sep (+/+)-*SCC+MIcyt, *Sep (-/-)*: *Sep (-/-)-*SCC+MIcyt, *Sep (-/-)* C2028S: *Sep (-/-)* C2028S-SCC+MIcyt. Bars, 10 µm. (D) A representative image of chromosome spreads showing that transferred bivalents from *Sep* null oocytes formed dyads after the first meiotic division. The antibody is shown in the panel (green). DNA was stained with DAPI (grey). Bar, 15µm.

### Different kinetics of REC8 cleavage along chromosome arms, at centromeres, and within peri-centromeric chromatin are conferred by SGOL2

The preceding experiment raises the possibility that some form of REC8 cleavage at the first meiotic division destroys peri-centromeric cohesin protection as well as kinetochore co-orientation. Because sister chromatid cohesion within centromeres presumably has a role in meiosis I kinetochore co-orientation, it is plausible to imagine that centromeric REC8 could be the key target. To explore this notion, we used super-resolution 3D structured illumination microscopy (3D-SIM) to observe in greater detail the location of meiotic specific kleisin REC8-containing cohesin as oocytes undergo the first meiotic division (Lee et al., 2003). We located centromeres using CREST (calcinosis, Raynaud’s phenomenon, esophageal dysmotility, sclerodactyly, telangiectasia) autoimmune serum and peri-centromeric chromatin using antibodies against histone H3 tri-methylated at lysine 9 (H3K9me3). Lastly, staining using antibodies specific for topoisomerase II revealed the two axes of each chromatid within peri-centromeric regions (Figs 4 and S4, Broccoli et al., 1990; Lee et al., 2011; Probst et al., 2007). Mice possess telocentromeric chromosomes. Their centromeres/kinetochores are therefore localized at the distal tips of each bivalent (Figs. 4A and S4A). Peri-centromeric chromatin, that is organized around axes associated with each sister chromatid, is located just proximal to these, with an axial length of 1.11 µm (Fig. S4A and B). During metaphase I, REC8 was detected between sister chromatids throughout bivalents, including peri-centromeric and centromeric regions. Despite their co-orientation, sister kinetochores are bisected by a zone of inter-chromatid REC8 when observed at this resolution. This is particularly clear when the bivalent is observed from the spindle pole to which it is attached (Fig. 4A MI-top). In other words, it is possible to detect a specific population of REC8 that is in an appropriate position to hold sister kinetochores together.

**Figure 4.**
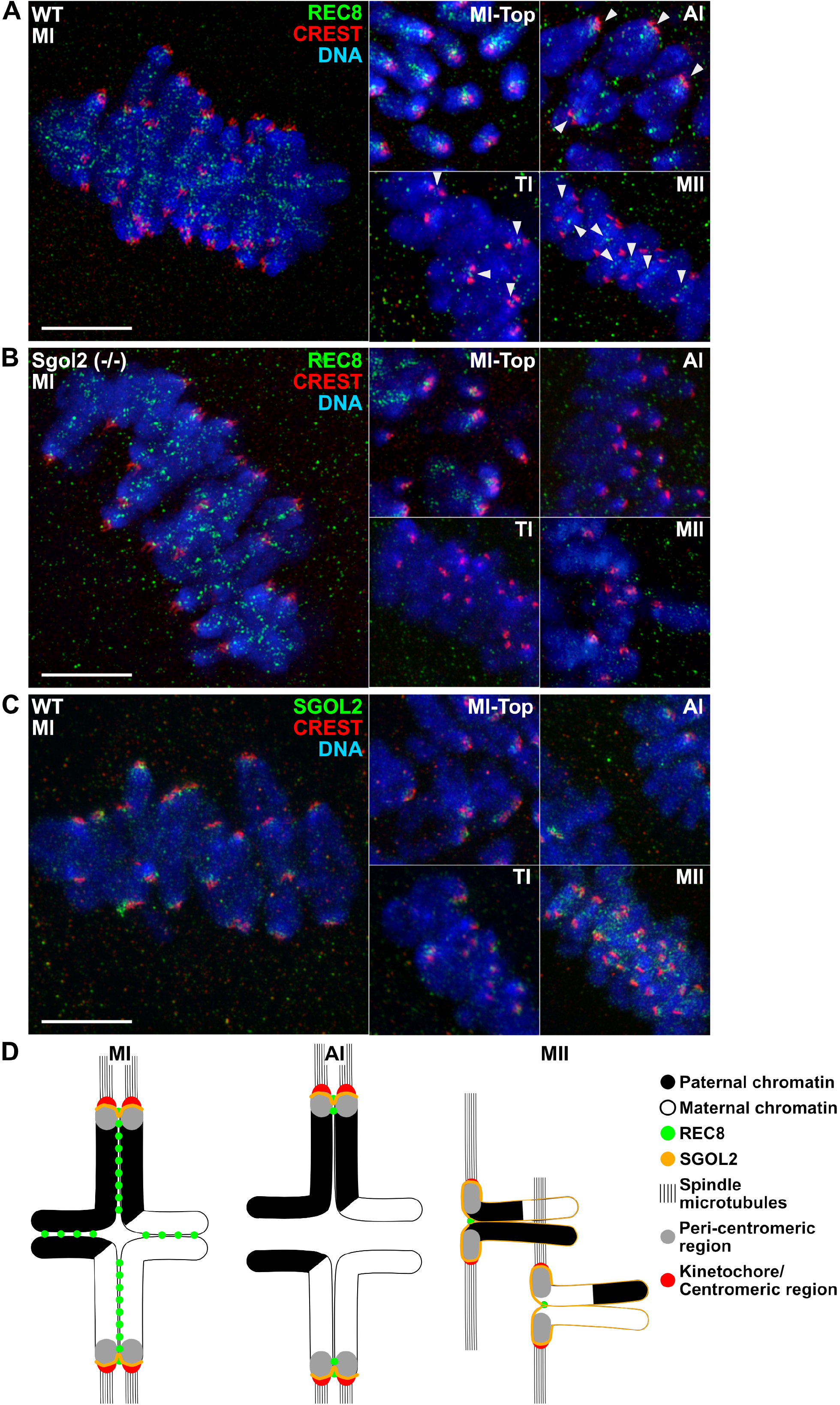
SGOL2 Determines the Dynamics of REC8 Cleavage of Arms, Centromeres, and Peri-centromeres during the First Meiotic Division. (A) Representative super-resolution 3D-SIM images showing REC8 localization in wild-type oocytes at MI, anaphase I (AI), telophase I (TI) and MII. The population of REC8 localized between sister kinetochores was observed from the top of spindle (MI-top). Arrowheads indicate REC8 in dyads at TI and MII. The antibodies are shown in the panel (green and red). DNA was stained with DAPI (blue). Bar, 5 µm. (B) Representative 3D-SIM images showing REC8 localization in *Sgol2 (-/-)* oocytes at MI, MI-top, AI, TI and MII, as in Figure 4A. Bar, 5 µm. (C) Representative 3D-SIM images showing SGOL2 localization in *wild*-type oocytes at MI, MI-top, AI, TI and MII, as in Figure 4A. Bar, 5 µm. (D) Schematic of results showing REC8 (green) and SGOL2 (orange) localization during MI-MII transition.

Soon after onset of the first anaphase (AI), REC8 disappears from chromosome arms, which leads to loss of sister chromatid cohesion at this location. This resolves the chiasmata and permits the traction of dyads towards the poles. Crucially, REC8 persists at this stage not only in peri-centromeric but also centromeric regions (Fig. 4A AI). Some also persists at both these locations until Telophase I (TI), during which a small amount remains sandwiched between sister kinetochores, which remain closely apposed. However, by the time cells enter MII, sister kinetochores are pulled several microns apart from each other and REC8, albeit in greatly reduced amounts, persists only in the peri-centromeric regions that hold the dyads together. These observations suggest that there are three populations of chromosomal REC8: arm, peri-centromeric, and centromeric. The arm population disappears at the onset of AI, the centromeric one only does so after TI, while the peri-centromeric population persists until the metaphase to anaphase division of meiosis II, albeit in greatly reduced amounts compared to meiosis I. All three populations of REC8 persist on bivalents when *Sep* (-/-) *C2028S* oocytes attempt the first meiotic division (Fig. S3A) but disappear simultaneously at the onset of AI in *Sgol2 (-/-)* oocytes (Fig. 4B). This suggests that SGOL2-PP2A protects centromeric as well as pericentromeric REC8 from separase. However, in the case of centromeric REC8, SGOL2 merely delays cleavage until after telophase. Surprisingly, SGOL2 accumulates to maximal levels not within peri-centromeric chromatin but just proximal to co-oriented kinetochores from metaphase to telophase during meiosis I (Fig. 4C) and only accumulates to higher levels within peri-centromeric chromatin during meiosis II. The high levels of SGLO2 next to kinetochores may help to explain how it delays cleavage of centromeric REC8. However, it is less clear why SGLO2 fails to protect this REC8 population when cells enter meiosis II but manages to protect at least some peri-centromeric REC8. The degree of protection cannot easily be explained by the pattern of SGOL2 accumulation.

### Centromeric REC8 facilitates sister kinetochore co-orientation

Separase destroys both co-orientation and protection of peri-centromeric cohesin as well as cleaving all chromosome arm and centromeric REC8 by the time cells enter meiosis II. Given this, either arm or centromeric REC8 cleavage could trigger loss of co-orientation and peri-centromeric protection. To address the role of centromeric REC8 cleavage, we injected oocytes whose *Rec8* contains Tobacco Etch Virus (TEV) protease cleavage sites (*Rec8-Tev)* with mRNAs that encode a fusion of the Cenp-C DNA binding sequence to the N-terminus of TEV (CCTEV) protease (Tachibana-Konwalski et al., 2010, 2013). By targeting CCTEV to centromeres, we aimed to cleave *Rec8-Tev* at centromeres but not elsewhere on chromosomes. The specificity of CCTEV was determined by checking the localization of REC8 on bivalents in *Rec8-Tev* oocytes 4-6 hours after injection of CCTEV mRNA at a concentration of 1 ng/µl (Fig. 5A and 5B). To prevent these oocytes from initiating anaphase, which would be accompanied by separase-mediated cleavage, they were injected at the GV stage with mRNAs for MAD2, which activates the SAC and prevents activation of the APC/C (Wassmann et al., 2003). To check whether the effects of CCTEV are due to its cleavage activity and not due to some adventitious side effect, we injected a set of control oocytes with a catalytically dead version, CCTEVC151A (Phan et al., 2002).

**Figure 5.**
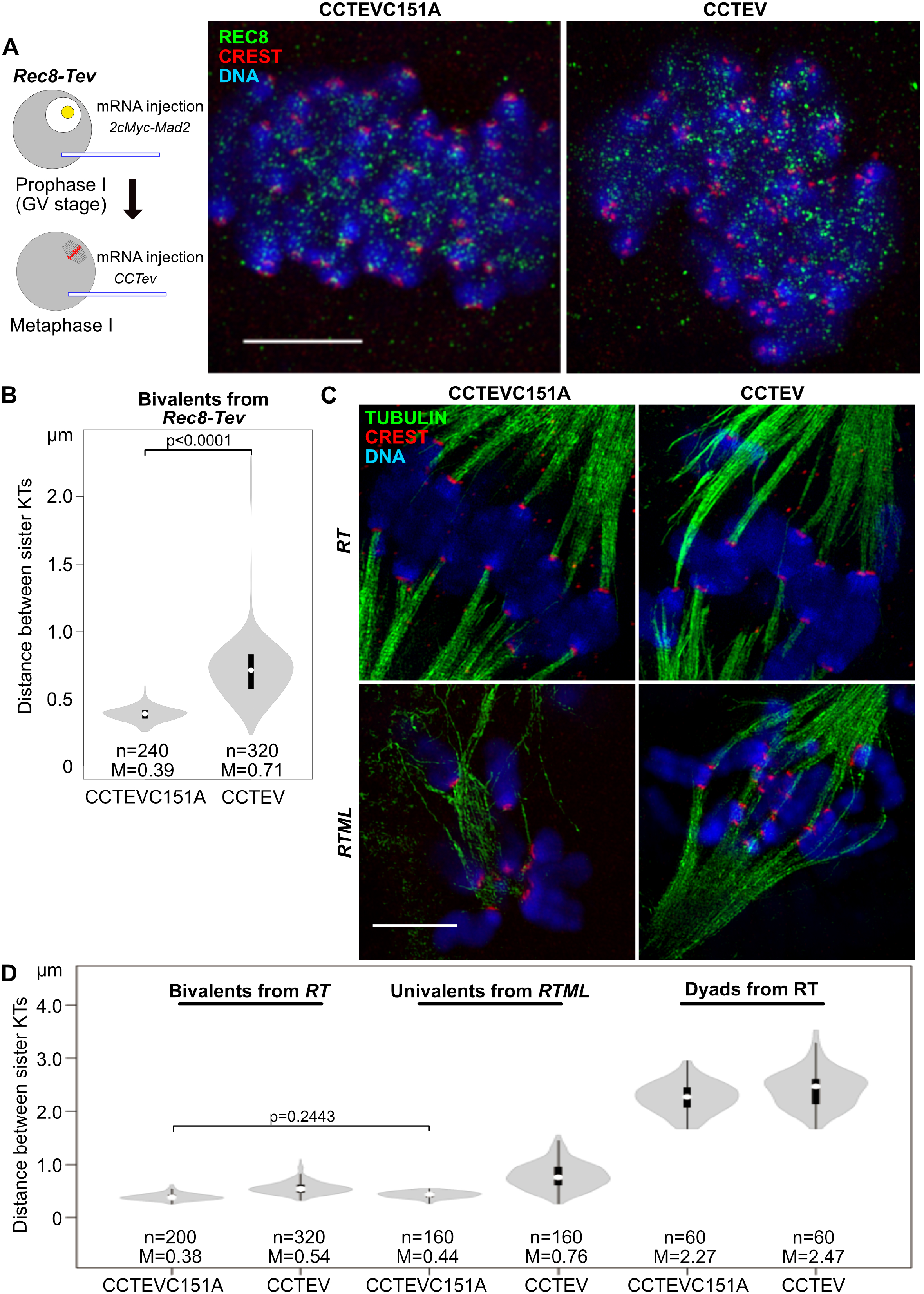
Centromeric Cohesin Is Necessary for Sister Kinetochore Co-orientation of Univalents at MI. (A) Left, Schematic of experiments showing that induction of centromeric specific cleavage in bivalents having TEV-cleavable REC8 (REC8-TEV) using TEV protease fused with CENP-C DNA binding domain (CCTEV). Right, A representative 3D-SIM image showing specific cleavage of REC8-TEV around centromeres 6 hours after induction of CCTEV expression. CCTEVC151A is a catalytic dead version of CCTEV. The antibodies are shown in the panel (green and red). DNA was stained with DAPI (blue). Bar, 5µm. (B) Distance between sister kinetochores (KTs) in bivalents 4-6 hours after cleavage of REC8-TEV around centromeres by CCTEV. White circles show the medians (M); box limits indicate the 25th and 75th percentiles as determined by R software; whiskers extend 1.5 times the interquartile range from the 25th and 75th percentiles; polygons represent density estimates of data and extend to extreme values. Two-tailed t-test was performed to test significance (p < 0.001). (C) A representative 3D-SIM image showing bi-orientation of sister kinetochores in univalents from *Rec8-Tev Mlh1* (-/-) (*RTML*) oocytes, which were lost centromeric cohesin as in Figure 5A. Bar, 5µm. (D) Distance between sister KTs in bivalents, univalents and dyads 4-6 hours after cleavage of REC8-TEV around centromeres by CCTEV. One-way ANOVA tests were performed to test significance (*p* < 0.001). Only non-significance was shown.

Six hours after mRNA injection, CCTEV but not CCTEVC151A caused the disappearance of REC8 from centromeres but not from chromosome arms, which was accompanied by a distinct moving apart of sister kinetochores (Fig. 5A and B). Sister kinetochore separation reached a median value of 0.71 µm, nearly twice as far as the 0.39 µm in control CCTEVC151A oocytes. Interestingly, sister kinetochore co-orientation persisted despite the clear loss of cohesion between sister centromeres (Fig. 5A and B). There are two potential explanations for this. One possibility is that CCTEV does not completely remove all centromeric REC8 and residual cohesin at this location is sufficient to support co-orientation. Alternatively, once co-orientation has been established, chiasmata ensure that tension stabilizes this state even when centromeric cohesin has been fully removed by CCTEV. To address the latter, we repeated the experiment using *Rec8-Tev Mlh1* (-/-) (*RTML*) oocytes, which are defective in recombination and contain univalent instead of bivalent chromosomes (Baker et al., 1996; Tachibana-Konwalski et al., 2010). The sister kinetochores of these univalents co-orient and because of this and because of the lack of chiasmata holding maternal and paternal kinetochore pairs together, their kinetochores cannot establish stable connections to microtubules. They instead move from pole to pole as they are pulled first one way and then another.

The univalents from *RTML* oocytes injected with CCTEVC151A mRNA behaved in a similar fashion but those injected with CCTEV underwent efficient bi-orientation and aligned in a stable manner on the spindle midplate (Fig. 5C). These observations confirm that CCTEV is indeed highly specific because cohesion persists along the inter-chromatid axes of univalents and thereby enables their stable bi-orientation. They also demonstrate that in the absence of chiasmata, cleavage of centromeric REC8 is sufficient to destroy co-orientation. Thus, centromeric REC8 has an important role in co-orientation but it is not necessary when chiasmata connect maternal and paternal sister kinetochore pairs. Whether a system analogous to the bundling of kinetochores by monopolin in yeast also has a role, especially in the absence of centromeric REC8, has not been addressed by our experiments.

The separation between sister kinetochores induced by cleavage of centromeric REC8 by CCTEV was slightly greater in oocytes containing univalents than those containing bivalents (Fig. 5D). This is not surprising as spindle forces pull apart the former but not the latter. More interestingly, we noticed that sister kinetochores separate less on bi-orienting univalents from *Rec8-Tev Mlh1* (-/-) (*RTML*) MI oocytes injected with CCTEV mRNA than on bi-orienting dyads from *Rec8-Tev (RT*) MII oocytes injected with either CCTEV or CCTEVC151A *mRNAs.* This suggests that separase removes more cohesin, either from centromeres or more likely from peri-centromeric chromatin, than does CCTEV. This implies that protection of peri-centromeric cohesin by SGOL2 is in fact only partial, a scenario that is consistent with the lower levels of REC8 associated with the peri-centromeric chromatin of dyads compared to bivalents (Fig.4A).

These experiments also revealed that separation between the sister kinetochores of dyads was slightly greater when *Rec8-Tev* meiosis II oocytes were injected with CCTEV than with *CCTEVC151A mRNAs* (Fig. 5D), raising the possibility that CCTEV may induce modest cleavage of peri-centromeric as well as centromeric cohesin. We therefore set out to establish more precisely the specificity of CCTEV using a functional assay. To do this, we transferred an SCC containing univalents from an *RTML* MI oocyte to an MII *RT* oocyte containing dyads and analysed the consequences of injecting either *CCTEV* or *CCTEVC151A mRNAs* (Fig. 6A). If CCTEV cleaves centromeric but not peri-centromeric REC8, then CCTEV should induce bi-orientation of the transferred univalents but not induce disjunction of the host dyads. In the oocytes injected with *CCTEVC151A mRNAs*, the transferred univalents moved back and forth between spindle poles while the host dyads aligned at the spindle midzone, forming a metaphase plate. In contrast, in oocytes injected with *CCTEV mRNAs*, both sets of chromosomes bi-oriented at the spindle zone and remained there for several hours (Fig. 6B, Movies 8 and 9), implying that CCTEV destroys within the very same cell sufficient centromeric REC8 to induce bi-orientation of univalents without adversely affecting the peri-centromeric cohesion holding dyads together. Chromosome spreads from these oocytes confirmed that CCTEV’s cleavage activity caused a modest increase in the separation of sister kinetochores associated with the univalents but not to the extent observed with dyads (Fig. 6C). In conclusion, by cleaving REC8, CCTEV triggers a change in the geometry of centromeric DNA, which induces univalent sister kinetochores to bi-orient, while leaving peri-centromeric cohesion more or less intact.

**Figure 6.**
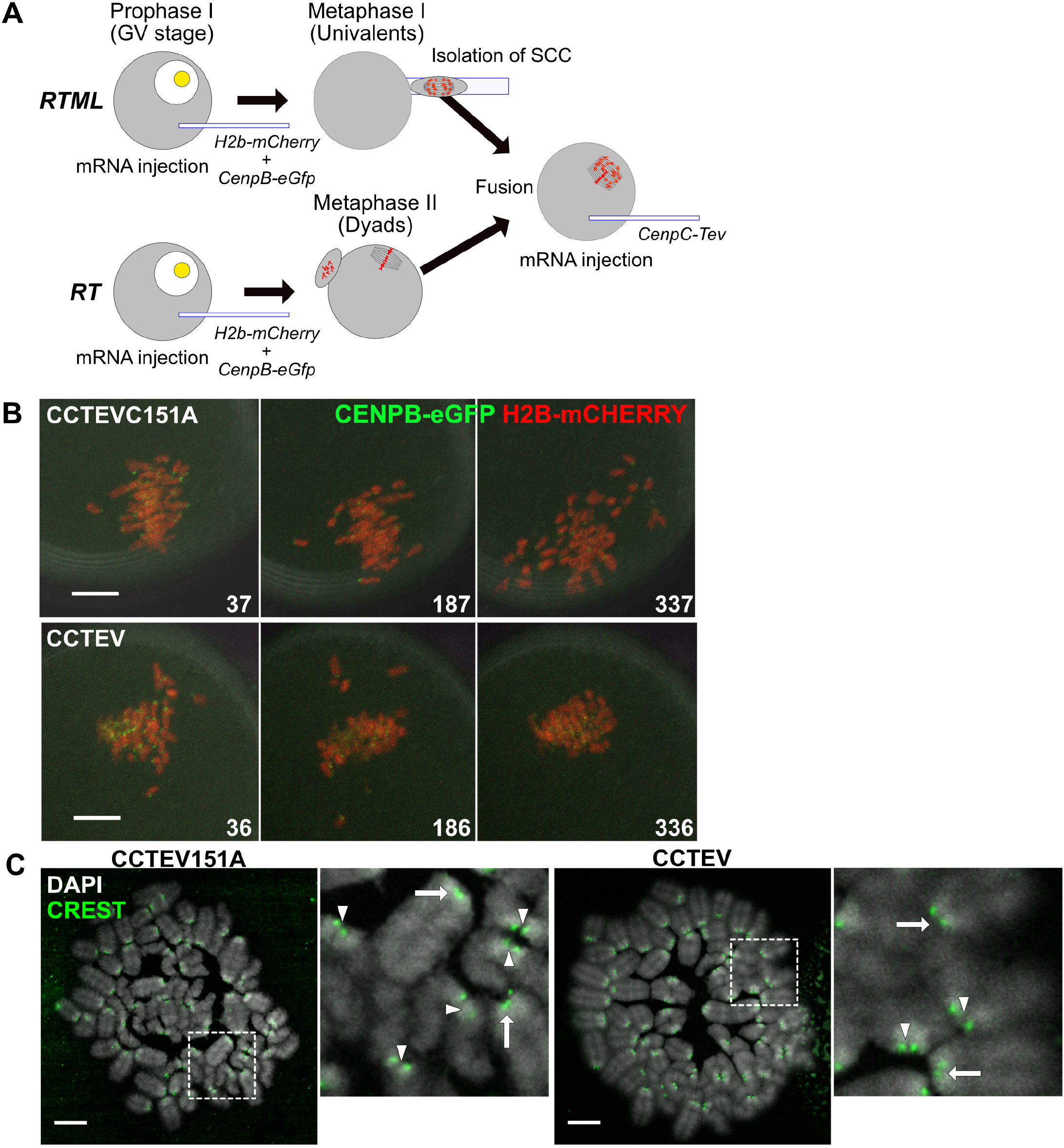
Specific Cleavage of TEV-Cleavable REC8 around Centromeres by CENPC-TEV. (A) Schematic of experiments showing an SCC having univalents (*Rec8-Tev Mlh1 (-/-): RTML*) is fused with an MII oocyte having dyads (*Rec8-Tev Mlh1(+/+): RT*) and then REC8-TEV around centromeres is cleaved by expression of CCTEV. (B) Representative stills from live cell imaging showing the induction of sister kinetochore bi-orientation in univalents by loss of centromeric cohesin under the condition that peri-centromeric cohesin in dyads was maintained. Chromosomes were labelled by H2B-mCHERRY (red) and kinetochores were visualized by CENPB-eGFP (green). Numbers indicate the time after induction of CCTEV expression (min). Bars, 10 µm. (C) A representative image of chromosome spread showing that specific cleavage of REC8-TEV around centromeres by CCTEV. Images beside the whole chromosome spread show 4-fold magnification of the regions indicated in the dash-lined boxes. Arrows indicate univalents and arrowheads indicate dyads. The antibody is shown in the panel (green). DNA was stained with DAPI (grey). Bars, 10 µm.

### Centromeric REC8 is necessary for efficient protection of peri-centromeric REC8

Because separase-mediated cleavage mediated by separase is required to destroy protection of peri-centromeric cohesin by SGOL2 as well as co-orientation, we next addressed whether centromeric REC8 is necessary for protection as well as co-orientation. To do this, we transferred to a wild-type oocyte an SCC containing univalents from *RTML* oocytes whose centromeric REC8 had been cleaved by CCTEV (Fig. 7A). To distinguish the transferred chromosomes from those of the host oocyte, the latter were isolated from B6D2F1 mice whose paternal chromosome 1 contains two centromeric regions separated by a peri-centromeric region (Mitchell et al., 1993), which associates with especially high levels of eGFP-CENPB. Fused oocytes containing univalents with centromeric REC8, namely those from *RTML* oocytes injected with *CCTEVC151A mRNAs*, failed to undergo the first meiotic division due to SAC activation (0/16, Fig. 7B and D) (Tachibana-Konwalski et al., 2013). In contrast, those lacking centromeric REC8, namely those from *RTML* oocytes injected with *CCTEV mRNAs*, did so with an average timing of 664±88 min (13/21, Fig. 7B and D) because bi-orientation of the univalents satisfied the SAC. Crucially, 75% of univalents were converted to individual chromatids at this division (Fig. 7E and F, *RTML*-CCTEV & Movies 10 and 11). Indeed, had this not occurred, the division would not have been possible. Thus, removing centromeric REC8 not only abolishes kinetochore co-orientation but also the protection of peri-centromeric REC8.

**Figure 7.**
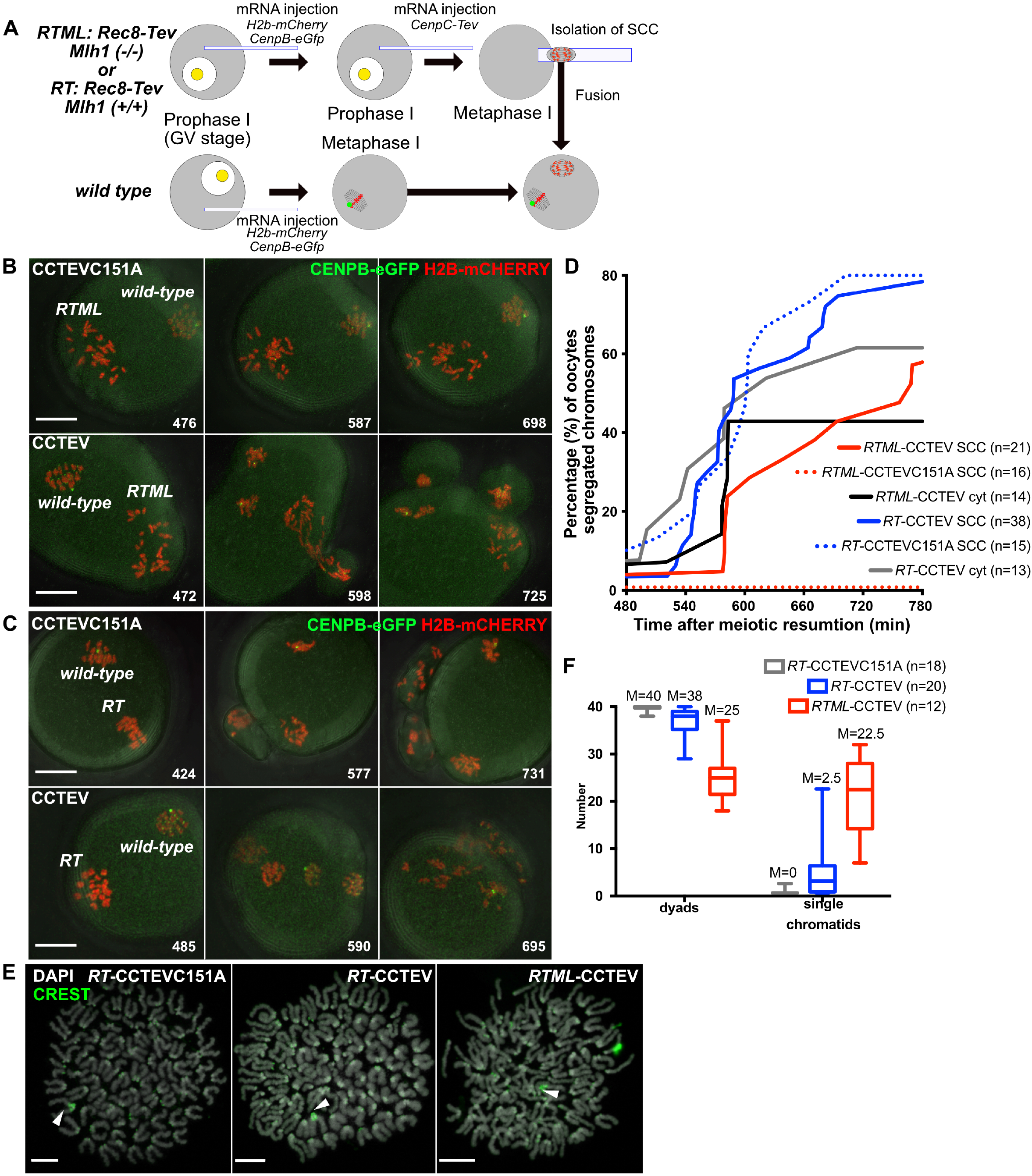
Centromeric Cohesin Is Necessary for Protecting Peri-centromeric Cohesin at the First Meiotic Division. (A) Schematic of experiments that demonstrate requirement of centromeric cohesin for protection of peri-centromeric cohesin. An SCC containing univalents (*Rec8-Tev Mlh1 (-/-): RTML*)/bivalents (*Rec8-Tev Mlh1 (+/+): RT*) without centromeric cohesin is fused with an MI oocyte (wild-type) from B6D2F1 mouse of which paternal chromosome 1 has two amplified centromeric regions separated by a peri-centromeric region thereby marked as a big bright signal by CENPB-eGFP. (B) Representative stills from live cell imaging showing that univalents (*RTML*) lacking centromeric cohesin converted into single chromatids after the first meiotic division as in Figure 6B. Numbers indicate the time after meiotic resumption of wild-type oocytes (min). Bars, 20 µm. (C) Representative stills from live cell imaging showing that some bivalents (RT) lacking centromeric cohesin converted to single chromatids as in Figure 6B. Bars, 20 µm. (D) Segregation timing of bivalents (*RT*)/univalents (*RTML*) lacking centromeric cohesin in an MI oocyte. Each group indicates wild-type oocytes fused with SCC (SCC) or cytoplasm (cyt) from *RT*/*RTML* after centromeric cohesin cleavage by CCTEV. (E) A representative image of chromosome spread showing that univalents lacking centromeric cohesin formed some single chromatids after the first meiotic division. Each panel indicates the formation of dyads or single chromatids in a wild-type MI oocyte fused with an SCC from an *RT/RTML* oocyte previously injected with CCTEV/*CCTEVC151A*. For confirming dyad formation of a paternal chromosome 1 from a wild-type oocyte (B6D2F1) having two amplified centromeric regions, cytokinesis was inhibited by cytochalasin D at 0.5 µg/ml. The antibody is shown in the panel (green). DNA was stained with DAPI (grey). Arrowheads indicated paternal chromosome 1 from wild type oocytes. Bars, 10 µm. (F) Number of dyad or single chromatid formation after the first meiotic division in a wild-type MI oocyte fused with an SCC from an *RT/RTML* oocyte previously injected with CCTEV/*CCTEVC151A*. Boxes show the median, 25th and 75th percentiles, and bars show the 10th and 90th percentiles. Median (M).

To address whether loss of protection is due to the bi-orientation of sister kinetochores caused by loss of centromeric REC8, we repeated the experiment, this time transferring bivalents from *RT* oocytes previously injected with *CCTEVC151A* or *CCTEV mRNAs*. In the case of former, all bivalents were converted to dyads when the oocytes underwent MI with an average timing of 586±64 min (12/16, Fig. 7C and D). However, 10% of bivalents whose centromeric REC8 had been removed by CCTEV were converted to individual chromatids not dyads when the oocytes underwent the first meiotic division with an average timing of 609±84 min. (31/39, Fig. 7C, D and F and Movies 12 and 13). Since none of the bivalents underwent bi-orientation in this experiment, it would appear that centromeric REC8 may have a direct role in conferring protection of peri-centromeric REC8. Interestingly, this premature loss of peri-centromeric cohesion was not accompanied by any change in the localization of SGLO2 and MEIKIN at MI (Fig. S5).

To determine whether loss of sister kinetochore co-orientation alone also contributes to deprotection of peri-centromeric REC8, we analysed the behavior of *Mlh1 (-/-)* oocytes with intact REC8. Though most of their sister kinetochores co-orient, we noticed that one or two pairs of sister kinetochores bi-orient (Fig. S6A). To test whether such chromosomes produce single chromatids when cells undergo the first meiotic division, we overcame the SAC-induced meiotic arrest caused by co-orientation of the majority of univalents, by expressing a dominant-negative version of the APC/C activator CDC20, CDC20R132A (Fig. S6C, Tachibana-Konwalski et al., 2013). CDC20 was used as an injection control and the first anaphase entrance was measured by monitoring degradation of SECURIN-eGFP (McGuinness et al., 2009). As expected, most univalents from *Mlh1 (-/-)* oocytes injected with *Cdc20R132A* mRNA were converted into dyads during a division whose average timing was 412±130 min (17/19). However, one or two univalents in each oocyte were instead converted to individual chromatids (Fig. S6D and E and Movies S6-S9), implying precocious loss of peri-centromeric cohesion. We presume that these univalents were those that had previously bi-oriented but, due to difficulties in tracking individual chromosomes throughout the division, were unable to establish this with any certainty. It is therefore possible that bi-orientation per se could also contribute to precocious loss of peri-centromeric cohesion. Crucially, this effect was not due to the abnormal nature of the division induced by *Cdc20R132A*, because bivalents from *Mlh1 (+/+)* oocytes invariably produced dyads even when their first meiotic division was accelerated. For example, oocytes injected with *Cdc20R132A* underwent the first meiotic division with an average timing of 321±83 min (14/18), which is substantially faster than those injected with *Cdc20* mRNA (5/8, 605±119 min) (Fig. S6B and C and Movies S6-S9).

### SGLO2 is dispensable for maintaining peri-centromeric cohesion during MII

How SGOL2 protects at least some peri-centromeric REC8 but merely delays cleavage of centromeric REC8 upon separase activation at the onset of anaphase remains an enigma since SGOL2 accumulates to much higher levels at centromeres during meiosis I than it does at peri-centromeres. It is nevertheless striking that SGOL2 does subsequently accumulate on peri-centromeric sequences during MII. To address whether its presence at this location is necessary to maintain the peri-centromeric cohesion holding dyads together prior to APC/C activation upon fertilization, we created a version of SGOL2 whose N-terminus was fused to eGFP and that contains 3xTEV recognition sites at cysteine 706 (SGOL2-TEV706)(Fig. S7A). Injection of *Sgol2-Tev706* mRNA at the GV stage had little or no effect on the frequency or timing of meiosis I of neither of *Sgol2 (+/+)* nor *Sgol2 (-/-)* oocytes (Fig. S7B and C and Movies S10-S14) but fully restored the retention of peri-centromeric cohesion in the latter (Fig. S7E). Crucially, injection at MI of mRNAs encoding TEV protease abolished this ability (Fig. S7B & D and Movies S10-S12), implying that cleavage around cysteine 706 inactivates SGLO2. In contrast, dyads created by injection of *Sgol2 (-/-)* oocytes with *Sgol2-Tev706* mRNA at the GV stage were unaffected when TEV mRNAs were instead injected during MII, implying that though required during anaphase I, SGOL2 is unnecessary for maintaining peri-centromeric cohesion once oocytes enter MII (Fig. S7D & E).

## Discussion

The work described in this paper addressed whether sister kinetochore co-orientation and shugohsin-mediated protection of peri-centromeric cohesin are conferred by the state of chromosomes or by physiological differences between the cytoplasm of meiosis I and meiosis II cells. Our observation that bivalents transplanted from meiosis I to meiosis II oocytes retain their meiosis I character while dyads transplanted from meiosis II to meiosis I oocytes retain their meiosis II character suggests that co-orientation and peri-centromeric cohesin protection are properties not of the cytoplasm but of the chromosome. Because of similar findings with grasshopper spermatocytes, (Nicklas, 1977; Paliulis and Nicklas, 2004), the chromosomal determination of kinetochore and pericentromeric chromatin behaviour appears to be a conserved feature, at least between insects and mammals.

An important advantage of the oocyte system is that it has been possible to address the molecular mechanism by which co-orientation and centromeric protection are destroyed during the transition from meiosis I to meiosis II using cytological and genetic manipulation. Our finding that cleavage activity mediated by separase is necessary suggests that it could be mediated through the cleavage of cohesin’s REC8 subunit. Crucially, no other change brought about through activation of the APC/C or changes in the activity of protein kinases during the transition from meiosis I to meiosis II is sufficient in the absence of separase-mediated cleavage.

Super resolution imaging with 3D-SIM revealed important differences in the kinetics of REC8’s cleavage by separase between chromosomal regions. While REC8 associated with chromosome arms disappears at the onset of anaphase I, an event that is necessary for chiasmata resolution, REC8 associated with centromeres persists until telophase or later. The improved resolution made possible by 3D-SIM enabled us to distinguish centromeric from peri-centromeric REC8, which in mice corresponds to approximately 1 µm in axial length. While all centromeric REC8 and some peri-centromeric REC8 disappeared at telophase I, a “protected fraction” of peri-centromeric REC8 persists until the onset of anaphase II. Crucially, SGOL2 is responsible for the differential behaviour as all three REC8 populations disappear with identical kinetics in *Sgol2* (-/-) oocytes, namely at the onset of anaphase I.

Loss of co-orientation and peri-centromeric protection as cells enter meiosis II could therefore be dependent on REC8 cleavage at any one of these three chromosomal locations. Though we were unable to block specifically cleavage at one but not the other two locations, we were able to test the effect of cleaving only centromeric REC8, which was achieved by injection into oocytes containing TEV-cleavable REC8 of mRNAs for a version of the TEV protease fused to CenpC’s DNA binding domain. Despite increasing the separation between sister kinetochores, cleavage of centromeric REC8 had little effect on the co-orientation of bivalents in MI. In contrast, it abolished co-orientation of sister kinetochores within univalents from *Mlh1* (-/-) oocytes, causing them to bi-orient and subsequently disjoin into individual chromatids at anaphase I. One explanation for this difference is that, once established microtubule-kinetochore attachments associated with bivalents are stable while those associated with univalents are unstable (because co-orientation prevents the establishment of tension) and loss of co-orientation may require the transient detachment of kinetochores from microtubules. We note that co-orientation is also more readily lost in the absence of chiasmata in the fission yeast, *S.pombe*, (Hirose et al., 2011; Sakuno et al., 2009; Sakuno et al, 2011; Yokobayashi & Watanabe, 2005). Though it is entirely plausible that centromeric cohesin co-orients kinetochores by holding together sister DNAs associated with kinetochore proteins, our experiments do not exclude the possibility that another type of activity, such as loop extrusion altering the topology of centromeric DNA (Davidson et al., 2019; Kim et al., 2019), might be involved.

Directed cleavage experiments similar to those described here have previously implicated centromeric cohesin in the co-orientation of sister kinetochores in *S.pombe*, suggesting that Rec8’s role may be evolutionarily conserved. However, in this case there was no clear evidence that Rec8 cleavage occurred specifically within centromeric as opposed to peri-centromeric chromatin (Yokobayashi & Watanabe, 2005). In budding yeasts with point centromeres, co-orientation depends on a monopolin complex that is thought to act by crosslinking not DNAs but kinetochore proteins (Corbett and Harrison, 2012; Furuyama and Biggins, 2007; Sarangapani et al., 2014). Whether centromeric cohesin is also required is not known. What is known, however is that co-orientation persists when the meiosis-specific kleisin subunit Rec8 is replaced by its mitotic equivalent Scc1 (Rabitsch et al., 2003; Sarangapani et al., 2014; Tóth et al., 2000), which is not the case in *S.pombe*.

Our experiments demonstrate that separase-mediated cleavage, presumably of REC8, also has a role in ablating protection by SGOL2 of peri-centromeric cohesin, whose removal triggers anaphase II. Two mysteries surround this phenomenon. The first concerns how protection in a form that lasts till the onset of the second meiotic division is only conferred on (some) peri-centromeric cohesion. There is a clear logic behind this. If as our work suggests co-orientation is conferred by centromeric cohesin, then bi-orientation of dyads during meiosis II requires the prior and complete removal of this cohesin population. Despite this logic, it remains unclear how SGOL2, which accumulates to far higher levels near centromeres than at peri-centromeric sequences, merely delays cleavage of centromeric REC8 until telophase I while protecting some peri-centromeric REC8 beyond this point in time. Protection of centromeric REC8 by SGOL2 must dissipate before separase is inactivated in telophase while that of peri-centromeric REC8 does not.

The second mystery concerns how protection within peri-centromeres does eventually dissipate by the time separase is re-activated at the onset of anaphase II. Cleavage of centromeric REC8 during meiosis I has a role in this dissipation as cleavage merely of centromeric REC8 induces not only bi-orientation of univalents but also their transformation into individual chromatids at anaphase I. However, it is also possible that limited cleavage of peri-centromeric REC8 also has a role in dissipation. In other words, protection by SGOL2 may require a critical concentration of peri-centromeric cohesin and partial cleavage of peri-centromeric REC8 during meiosis I may lower its level below this critical level. Unfortunately, our experiments did not establish with any certainty whether the loss of peri-centromeric protection caused by cleavage of centromeric REC8 is a direct consequence of REC8 cleavage or an indirect consequence of the bi-orientation of sister kinetochores induced by this cleavage. What is clear is that bi-orientation does not per se remove SGLO2 from peri-centromeric sequences, as has been proposed (Gómez et al., 2007; Lee et al., 2008), because SGOL2 accumulates at this location to high levels while sister kinetochores bi-orient during meiosis II. Interestingly, this phenomenon is irrelevant for maintaining the peri-centromeric cohesion holding dyads together during meiosis II, presumably because separase is insufficiently active until fertilization re-activates the APC/C. Indeed, our demonstration that SGOL2 can be inactivated through proteolytic cleavage during meiosis II without functional consequences is inconsistent with the suggestion that the dissipation of protection after meiosis I is mediated by the binding of a conserved histone chaperone SET/TAF-1b specifically at peri-centromeric region during meiosis II (Chambon et al., 2013; Wassmann, 2013). Spo13/Moa1/MEIKIN are meiosis-specific proteins from budding yeast, fission, and mammals that share the ability to bind to Polo-like kinases and have roles in both co-orientation of sister kinetochores and protection of cohesion (Galander et al., 2019; Katis et al., 2004a; Kim et al., 2015), raising the possibility that Spo13-like proteins may regulate the activity of centromeric cohesin.

## Materials and Methods

### Animal maintenance, handling and care, and mouse strains

All experimental procedures were approved by the University of Oxford local ethical review committee and licensed by the Home Office under the Animal (Scientific Procedures) Act 1986. For collection of wild-type oocytes, B6D2F1 mice were purchased from the Charles River or crossed in house between the C57B6/J female and DBA2 male mouse strains. *Zp3Cre Espl1(f/f)* (*separase (-/-)*) mice were produced by crossing *Espl1(f/f)* females and *Zp3Cre* heterozygous males (Kudo et al., 2006). *Rec8-Tev* mice were maintained as homozygous (Tachibana-Konwalski et al., 2010). *Mlh1* (-/-) *Rec8-Tev* mice were obtained by crossing *Mlh1* (+/-) *Rec8-Tev* mice.

### Modification of *CenpC-Tev* and *CenpC-TevC151A* constructs

The original plasmids for *CenpC-Tev/TevC151A* constructs, pRNA-*CenpC-mCherry-Tev*/*TevC151A* (kind gifts from K. Tachibana, IMBA, Austria) contain *mCherry* sequence between the sequences of *CenpC*-DNA binding domain and *Tev*. We improved the efficiency in kinetochore targeting of TEV by excising *mCherry* sequence and used the constructs, pRNA-*CenpC-Tev*/*TevC151A* in this study.

### mRNA synthesis and microinjection

Capped mRNA constructs with a poly-A tail were transcribed using an mMESSAGEmMACHINE kit containing the appropriate RNA polymerase (Ambion). Following Turbo DNase I digestion for 15 min at 37°C, RNA was purified by using RNeasy mini kit (Qiagen), and resuspended in nuclease-free H_2_O. Each RNA was aliquoted and stored at −80°C until use. We used following plasmid cDNA encoding *CenpB-eGfp, CenpC-Tev, CenpC-TevC151A, Cdc20, Cdc20R132A, eGfp-CenpC, eGfp-Sgol2, eGfp-Sgol2-Tev706, H2B-mCherry, Securin-eGfp, Espl1 (separase), separaseC2028S. In vitro*-transcribed RNAs of 5 pl were microinjected under inverted microscope (Leica DM IRB) equipped with a micromanipulator (Narishige) and a pressure injector (WPI PV830 PicoPump) to the oocytes at following concentrations. *CenpB-eGfp* 100 ng/µl, *CenpC-Tev* 1 ng/µl, *CenpC-TevC151A* 1 ng/µl*, Cdc20* 100 ng/µl*, Cdc20R132A* 100 ng/µl*, eGfp-CenpC* 500 ng/µl, *eGfp-Sgol2* 250 ng/µl*, eGfp-Sgol2-Tev706* 250 ng/µl*, mCherry-H2B* 200 ng/µl, *Securin-eGfp* 500 ng/µl*, separase* 100 ng/µl*, separaseC2028S* 100 ng/µl.

### Spindle isolation and fusion

Oocytes at the germinal vesicle (GV)-stage were collected from ovaries of female mice at 8−12 weeks of age at 44−48 hours after injection with 7.5 IU equine chronic gonadotropin (eCG; Intervet). Fully grown GV-oocytes were released from ovarian follicles by puncture with needles in M2 (Sigma) supplemented with 200 μM 3-isobutyl-1-methylxanthine (IBMX; Sigma), and their cumulus cells were removed by gentle pipetting. GV-oocytes were cultured for 1 hour in M16 containing 200 μM IBMX at 37°C under an atmosphere of 5% CO_2_, and then microinjected with mRNAs transcribed *invitro*. Injected oocytes were cultured for 1-2 hours in M16 containing 200 μM IBMX for expression of mRNAs, followed by the transfer into M16 for resumption of meiosis. After 4-6 hours from meiotic resumption, isolation of spindle was performed in M2 with 10-15 μg/ml cytochalasin D (Sigma) under an inverted microscope equipped with a micromanipulator and a PIEZO drive (Prime Tech PMAS-CT150). Firstly, an oocyte was positioned using a holding pipette so that the spindle was situated close to the 2 o’clock position. The zona pellucida next to a spindle was drilled with piezo pulses, an enucleation pipette was inserted through the opening hole, and a spindle surrounded by membrane was aspirated. Secondly, an enucleation pipette containing a spindle was move into the drop of HVJ-E extract (Ishihara Sangyo Kaisha) and HVJ-E extract with the double volume of spindle was aspirated. Finally, a spindle and HVJ-E extract were placed into the perivitelline space of the host oocyte. This construct was washed by M2 and transferred to M16, and incubated at 37°C in 5% CO_2_ for 30 min until fusion occurred.

### Parthenogenetic activation

For parthenogenetic activation, oocytes 1 hour after fusion were artificially activated by treating them with activation medium, consisted of modified Tris-buffered medium (20 mM Tris, 113 mM NaCl, 3 mM KCl, 11 mM glucose, 5 mM Na pyruvate, and 2 mg/ml BSA) with 10 mM SrCl_2_ and 5 μg/ml cytochalasin B, for 4-6 hours.

### Live cell imaging

Following fusion or artificial activation, oocytes were transferred to 5-10 μl drops of M16 or activation medium on a glass-bottomed dish and placed in an incubator (Zeiss) at 37°C under 5% CO2 in air. Observation was performed under an inverted confocal microscope (Zeiss LSM780) equipped with a C-Apochromat 63×/1.2 water immersion objective with 3D multilocation tracking macro, which was kindly provided by J Ellenberg (EMBL, Germany, Rabut and Ellenberg, 2004). We imaged 26 z-confocal sections every 2.0 μm of 512 × 512 pixel xy images at 15 min intervals.

### Chromosome spread

The zona pellucida of oocytes or parthenotes was removed by treatment with 5 mg/ml protease in M2; then, the oocytes/parthenotes were placed on a glass slide that had been dipped in a solution of 1% PFA in distilled water, pH 9.2, containing 0.15% Triton X-100 and 3 mM dithiothreitol. Following overnight fixation in a humid chamber, the slide was dried for 30 min at room temperature. The samples were washed in PBS and then incubated with anti-centromere protein (Antibodies Inc. #15-234-0001, 1:100) overnight at 4°C. After washing with PBS, the samples were incubated with Alexa-Fluor-488-conjugated donkey anti-human IgG for 1.5 hours at room temperature and mounted with Vectashield containing DAPI (Vector Laboratories).

### Whole-mount immunofluorescence

Basically, cultured oocytes were fixed in 2% paraformaldehyde (PFA) in PBS containing 0.1% Triton X-100 for 30 min. Only for microtubule staining, cultured oocytes were preincubated with ice-cold M2 for 10 min and proceeded to fixation as described above. After permeabilization with 0.1% Triton X-100 in PBS overnight at 4°C, oocytes were incubated with primary antibodies overnight at 4°C. Following three washes with PBS containing 0.1% polyvinyl alcohol (PVA), Alexa-Fluor-labelled secondary antibodies (Invitrogen) were used for the detection of signals and DNA was counterstained with 14.3 μM 4′,6-diamidino-2-phenylindole, dihydrochloride (DAPI). The following primary antibodies were used: anti-REC8 (a kind gift from J. Lee, Kobe University, Japan; 1:100), anti-centromere protein (1:100), anti-SGOL2, anti-MEIKIN (kind gifts from Y. Watanabe, University of Tokyo, Japan; 1:100 and 1:100, respectively), anti-TOPOII (Abcam, #ab109524, 1:100), anti-histone H3 tri-methylated at lysine 9 (H3K9me3; Abcam, #ab8898, 1:250-500) and anti-alpha TUBULIN (Sigma, #T9026, 1:250-500). Samples were mounted on a slide with Vectashield, covered with a No.1.5H (170 μm ± 5 μm) coverslip (Marienfield) and were imaged using a confocal laser scanning microscopy.

### 3D structured illumination microscopy (3D-SIM)

3D-SIM was performed on a DeltaVision OMX V3 Blaze system (GE Healthcare) equipped with sCMOS cameras (PCO), and 405, 488 and 593nm lasers, using a 60x NA 1.3 silicone immersion objective lens (Olympus). To minimize artefacts due to spherical aberration when imaging entire oocytes of increased height (~10-15 µm) and at extended depth (~20-30 µm), the samples were mounted on a microscope slide with 63% Vectashield diluted in PBS to match the refractive index of the silicone immersion medium (RI=1.40), and then covered with a No.1.5H coverslip. The correction collar of the objective was before adjusted to obtain a symmetrical point spread function when imaging green beads at 488 nm excitation. Raw data was acquired with a z-distance of 125 nm and with 15 raw images per plane (5 phases, 3 angles). The raw data was reconstructed with SoftWoRx 6.2 (GE Healthcare) using channel-specifically measured optical transfer functions (OTFs) generated from ~170nm diameter blue PS-Speck beads (ThermoFisher) and 100 nm diameter green and red FluoSphere beads (ThermoFisher), respectively, and Wiener filter setting 0.0040. Lateral color channel alignment was performed using a special image registration slide and algorithm provided by GE Healthcare. Correct 3D alignment was confirmed and refined in z by a custom test sample with two layers of 0.2 μm diameter TetraSpeck beads (ThermoFisher). The full-scale 32-bit reconstructed data was thresholded for each channel to the stack modal grey value (representing the centre of the background intensity level) and converted to 16-bit composite tif-stacks using an in-house script in ImageJ (http://rsbweb.nih.gov/ij) before further processing. All 3D-SIM data was evaluated via SIMcheck, an open-source ImageJ plugin to assess SIM image quality.

### Measure the distance between sister kinetochores and axis length of peri-centromeric regions

For measuring sister kinetochore distance, we used 3D-SIM images of CREST signals which have been reconstructed and thresholded. Using plugins in ImageJ, CREST signals were segmented by 3D segmentation and the sister kinetochore distance was measured by distance function in 3D manager. Axis lengths of peri-centromeric regions were measured using images of H3K9me3 signals. Images were thresholded, and signals were segmented by 3D segmentation. Axial lengths were measured by 3D measure function (Fit Ellipse) in 3D manager.

### Statistical Analysis

For statistical analysis, we used Two-tailed t-test (Fig. 5B) or One-way ANOVA tests (Fig. 5D) to compare kinetochore distance in bivalents and univalents before and after centromeric REC8 cleavage.

## Acknowledgement

We thank, J. Lee for anti-REC8 antibody, and Y. Watanabe for anti-SGOL2 and anti-MEIKIN antbodies, K. Tachibana for the pRNA*-CenpC-mCherry-Tev/TevC151A* plasmids, J. Ellenberg for cell tracking macro and B. Akiyoshi, J. Lee, T.S. Kitajima, and members of the Nasmyth and Turner lab for their critical input. Imaging was performed at the Micron Oxford Advanced Bioimaging Unit funded by a Wellcome Trust Strategic Award (091911 and 107457/Z/15/Z). This work was supported in part by a Grant-in-Aid from MEXT (to S.O.), by the Hakubi Center for Advanced Research (to S.O.), Wellcome Trust Programme Grants Ref 091859/Z/10/Z & Ref 107935/Z/15/Z (to K.N) and Cancer Research UK Programme Grant Ref C573/A12386 (to K.N).

## Author Contributions

S.O. designed, performed, and analysed all experiments, mounted figures, and wrote the manuscript; A.R. designed and performed some experiments and wrote the manuscript; J.G. and J.M performed experimental procedures; L.S. supervised SIM microscopy and contributed to data interpretation and writing the manuscript; K.N. supervised the work, designed and analysed experiments, and wrote the manuscript.

