## Supplemental Figure Legends for "Loss of Sister Kinetochore Co-orientation and Peri-centromeric Cohesin Protection after Meiosis I Depends on Cleavage of Centromeric REC8"

**Supplemental Figure 1**	**Either Sister Kinetochore Bi-orientation or De-protection of Cohesion Is Conferred by the Nature of Dyads Itself rather than the Surrounding Factors Coming from Cytoplasm**

(A) Representative stills from live cell imaging of dyad segregation in an MI oocyte without fusion procedure (MI) and in that fused with an MII cytoplast (MIIcyt+MI). Chromosomes were labeled by H2B-mCHERRY (red) and kinetochores were visualized by eGFP-CENPC (green). Numbers indicate the time after meiotic resumption (min). Bars, 10 µm.
(B) Timing of dyad segregation. SCC containing dyads from MII was fused with an oocyte at the germinal vesicle (GV) stage (MII-SCC+GV). As a technical control, MII cytoplasm was fused with a GV oocyte (MIIcyt+GV). n, the numbers of oocytes measured in more than three independent experiments.
(C) Representative stills from live cell imaging of dyad segregation in MII-SCC+GV and in MIIcyt+MI as in Supplemental Figure 1A. Bars, 10 µm.
(D) A representative still from live cell imaging that showing dyad biorientation in MII-SCC+GV as in Supplemental Figure 1A. Arrow heads indicate bi-orientated dyads and arrows indicate dyads failed to associate with a spindle. Bar, 10 µm.

**Supplemental Figure 2**	**Determination of Protection of Cohesion or Sister Kinetochore Co-orientation Is Granted by Bivalents rather than the Factors from Cytoplasm**

Representative stills from live cell imaging of bivalent segregation in an MII oocyte without fusion procedure (MII) and an MII oocyte fused with an MI cytoplast (MIcyt+MII) as in Supplemental Figure 1A. Bars, 10µm.

**Supplemental Figure 3	Dyads that Have Undergone Two Successive Meiotic Divisions Are Converted to Single Chromatids after Artificial Strontium Activation.**

(A) Representative images showing bivalents from a *Zp3Cre* *separase* (f/f) oocytes (*Sep* (-/-)) at MI, whose meiotic arrest defects were rescued by expression of separase at the GV stage (*Sep* (-/-)+SEP), and *Sep* (-/-) oocytes at MII induced only cytokinesis by expression of catalytic-dead version of separase C2028S (*Sep* null C2028S). The antibodies shown in the panels (green and red). DNA was stained with DAPI (grey). Bars, 10µm.
(B) Segregation timing after artificial strontium activation in dyads from *Sep* (-/-) oocytes. *Sep* (-/-)+SEP-SCC+MIcyt: an SCC from a *Sep* (-/-)+SEP oocyte was fused with an MI cytoplast from wild-type, *Sep* (-/-) C2028S-SCC+MIcyt: an SCC from a *Sep* (-/-) oocyte induced only cytokinesis by expression of catalytic-dead version of separase C2028S was fused with an MI cytoplast from wild-type. In both groups, dyads were prepared by induction of MI-MII transition after fusion. n, the numbers of oocytes measured in more than two independent experiments.
(C) Representative stills from live cell imaging showing that dyads that have successfully undergone three successive divisions after artificial activation as in Supplemental Figure 1A. Numbers indicate the time after artificial activation (min). *Sep* (-/-)+SEP: *Sep* (-/-)+SEP-SCC+MIcyt, *Sep* (-/-) C2028S: *Sep* (-/-) C2028S-SCC+MIcyt. Bars, 10 µm.
(D) A representative image of chromosome spread showing the formation of single chromatids from dyads that have undergone three successive divisions after artificial activation. The antibody is shown in the panel (green). DNA was stained with DAPI (grey). Bar 15 µm.

**Supplemental Figure 4	REC8 Localization at Centromeric, Peri-centromeric and Arm Regions in Bivalents (MI), and at the Distal Side of Peri-centromeric Regions in Dyads (MII)**

(A) Centromeric regions were marked by CREST and peri-centromeric regions were marked by histone H3 tri-methylated at lysine 9 (H3K9me3) (Top) or topoisomerase II (TOPOII) (Bottom). The antibodies are shown in the panels (green, red and grey). DNA was stained with DAPI (blue). Bars, 5 µm.
(B) The axis length of peri-centromeric regions in mouse oocytes at MI. White circles show the medians (M); box limits indicate the 25th and 75th percentiles as determined by R software; whiskers extend 1.5 times the interquartile range from the 25th and 75th percentiles; polygons represent density estimates of data and extend to extreme values.

**Supplemental Figure 5	Disappearance of Centromeric REC8 Does Not Alter Localization of SGOL2 at MI**

Left, Schematic of experiments showing that the induction of specific cleavage of centromeric REC8 by CCTEV. Right, A representative 3D-SIM image showing that retained localization of SGOL2 and MEIKIN at centromeric regions 6 hours after induction of CCTEV expression. CCTEVC151A is a catalytic dead version of CCTEV. Insets show 1.5-fold magnification of the regions indicated in the dash-lined boxes. The antibodies are shown in the panel (green and red). DNA was stained with DAPI (blue). Bars, 5 µm.

**Supplemental Figure 6	Loss of Co-orientation of Sister Kinetochores Triggers Deprotection of Peri-centromeric REC8 during MI-MII Transition**

(A) A representative 3D-SIM image showing bi-orientation of sister kinetochores in two univalent (arrow heads) from *Mlh1* (-/-) oocytes. The antibodies are shown in the panel (green and red). DNA was stained with DAPI (blue). Bar 5µm.
(B) Segregation timing of bivalents/univalents after expression of CDC20R132A in an MI oocyte. n, the numbers of oocytes measured in more than two independent experiments.
(C) Schematic of experiments (left) and representative stills (right) showing the induction of first meiotic division by inhibition of the spindle check point using expression of CDC20R132A in *Mlh1*(-/-) oocytes. Chromosomes were labeled by H2B-mCHERRY (red) and transition of meiosis I-II was visualized by SECURIN-eGFP (green), whose destruction indicates first anaphase entry and re-accumulation shows entry to meiosis II. Bars, 10 µm.
(D) A representative image of chromosome spread showing formation of dyads from univalents. The antibody is shown in the panel (green). DNA was stained with DAPI (grey). Bar, 10 µm.
(E) Number of dyad or single chromatid formation after the first meiotic division in *Mlh1*(-/-) oocytes injected with *CDC20R132A* mRNA. Boxes show the median, 25th and 75th percentiles, and bars show the 10th and 90th percentiles. Median (M).

**Supplemental Figure 7**	**SGOL2 in meiosis II Is Not Required for Protection of Cohesin in Dyads**

(A) Insertion site of 3xTEV-recognition sequence in *Mus musculus* SGOL2 at cysteine 706. Each colored box shows a binding region of indicated protein or conjugated eGFP at N terminus.
(B) Representative stills from live cell imaging showing that the defect of single chromatid formation in *Sgol2 (-/-)* oocytes was rescued by expression of TEV-cleavable SGOL2, SGOL2-TEV706 (green, middle), but not by expression of eGFP-CENPC (green, top). Expression of TEV at MI induced single-chromatid formation in *Sgol2 (-/-)* oocytes rescued by expression of SGOL2-TEV706 (green, bottom). Chromosomes were labeled by H2B-mCHERRY (red). Labels on each left top panel indicate a type of injected *Tev* mRNA at MI. Numbers indicate the time after meiotic resumption (min). Bars, 10 µm.
(C) Segregation timing of bivalents in *Sgol2 (-/-)* oocytes after expression of SGOL2-TEV706. After *Sgol2-Tev706* mRNA injection at the GV stage, chromosome segregation occurred at the average timing of 602±96 min and 595±118 min in *Sgol2 (+/+)* and *Sgol2 (-/-)* oocytes, respectively; 75% (58/77) of *Sgol2 (+/+)* and 76% (102/134) of *Sgol2 (-/-)* oocytes were extruded a polar body. All bivalents in *Sgol2 (-/-)* oocytes that had injected *Sgol2-Tev706* mRNA (300 ng/µl) at the GV stage and *TEVC151A* mRNA (300 ng/µl) at MI, respectively, segregated to dyads at the rate of 50% (12/24). In contrast, the frequency of chromosome segregation in *Sgol2 (-/-)* oocytes that had injected *Sgol2-Tev706* mRNA at the GV stage and, subsequently, *TEV* mRNA at MI, was at 47% (7/15) and all bivalents from these oocytes segregated to single chromatids. Numbers indicate the time after meiotic resumption (min). n, the numbers of oocytes measured in more than two independent experiments.
(D) Representative stills from live cell imaging showing the maintenance of dyads even after cleavage of SGOL2-TEV706 at MII. Labels on each left top panel indicate a type of injected *Tev* mRNA at MII. Numbers indicate the time after induction of TEV expression (min). Bars, 10 µm.
(E) A representative image of chromosome spread showing that the maintenance of dyads even after destruction of SGOL2 function at MII. The antibodies are shown in the panel (green and red). DNA was stained with DAPI (grey). Bar, 10 µm.
