## Supplementary figures and images for "Loss of Sister Kinetochore Co-orientation and Peri-centromeric Cohesin Protection after Meiosis I Depends on Cleavage of Centromeric REC8"

### Supplemental Figures

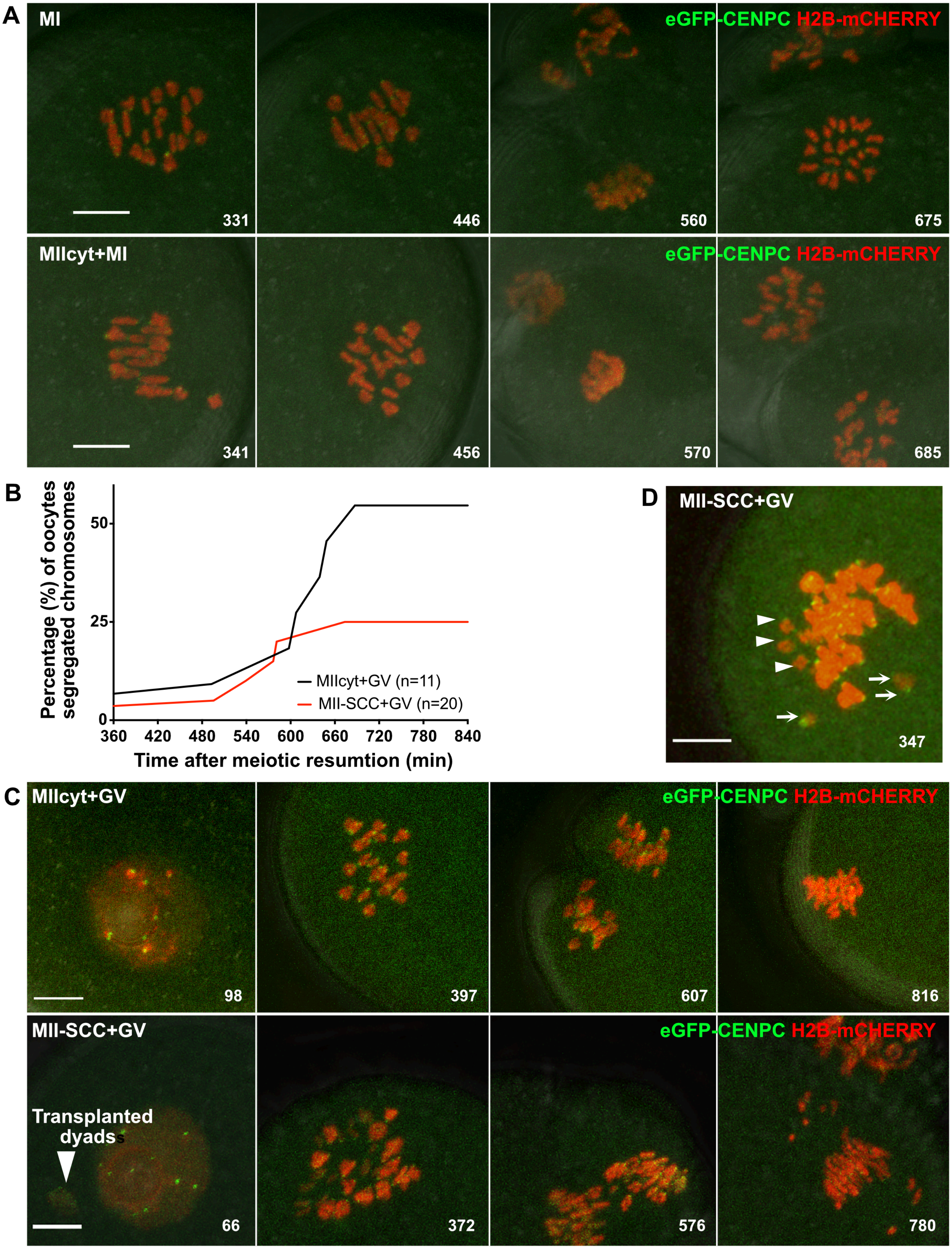

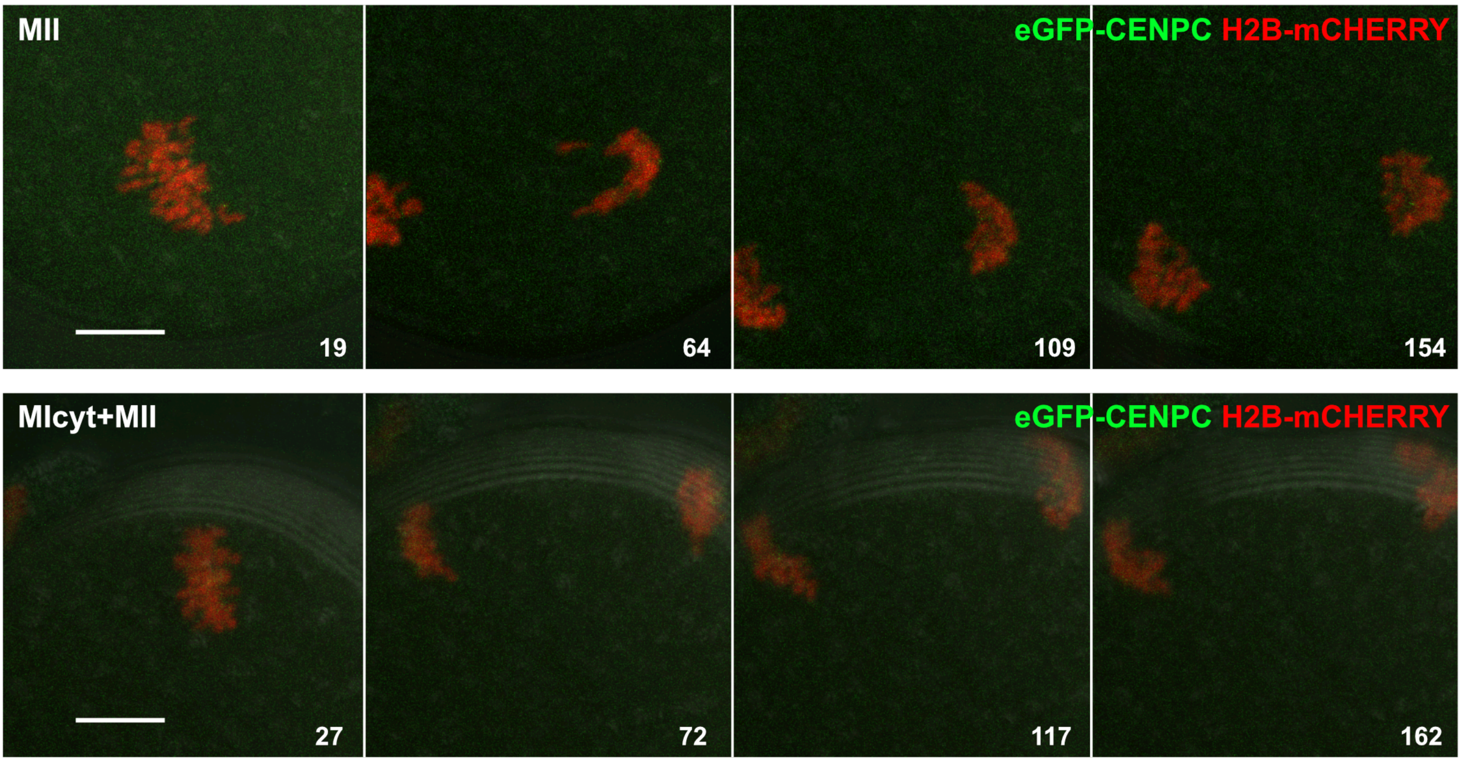

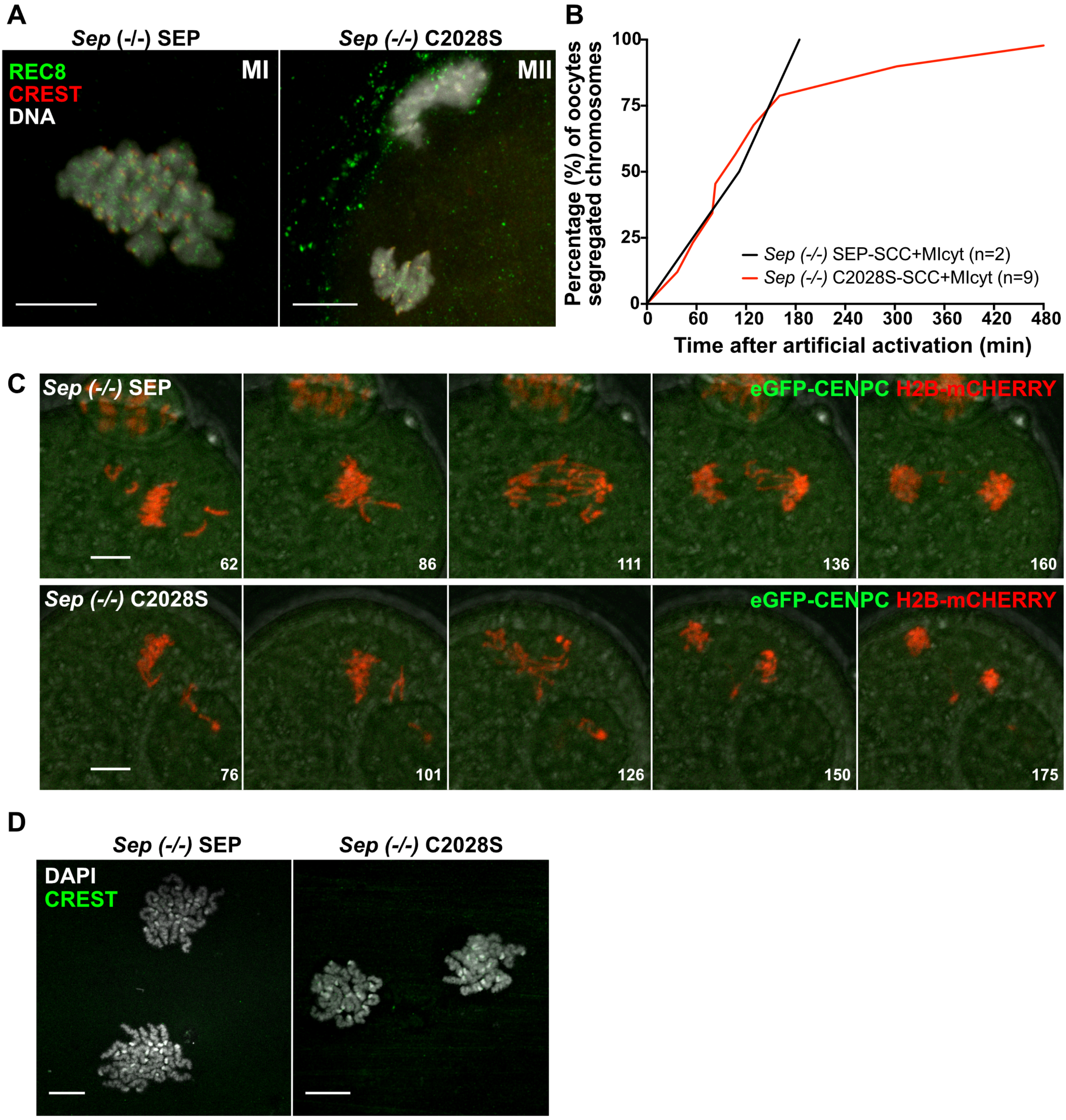

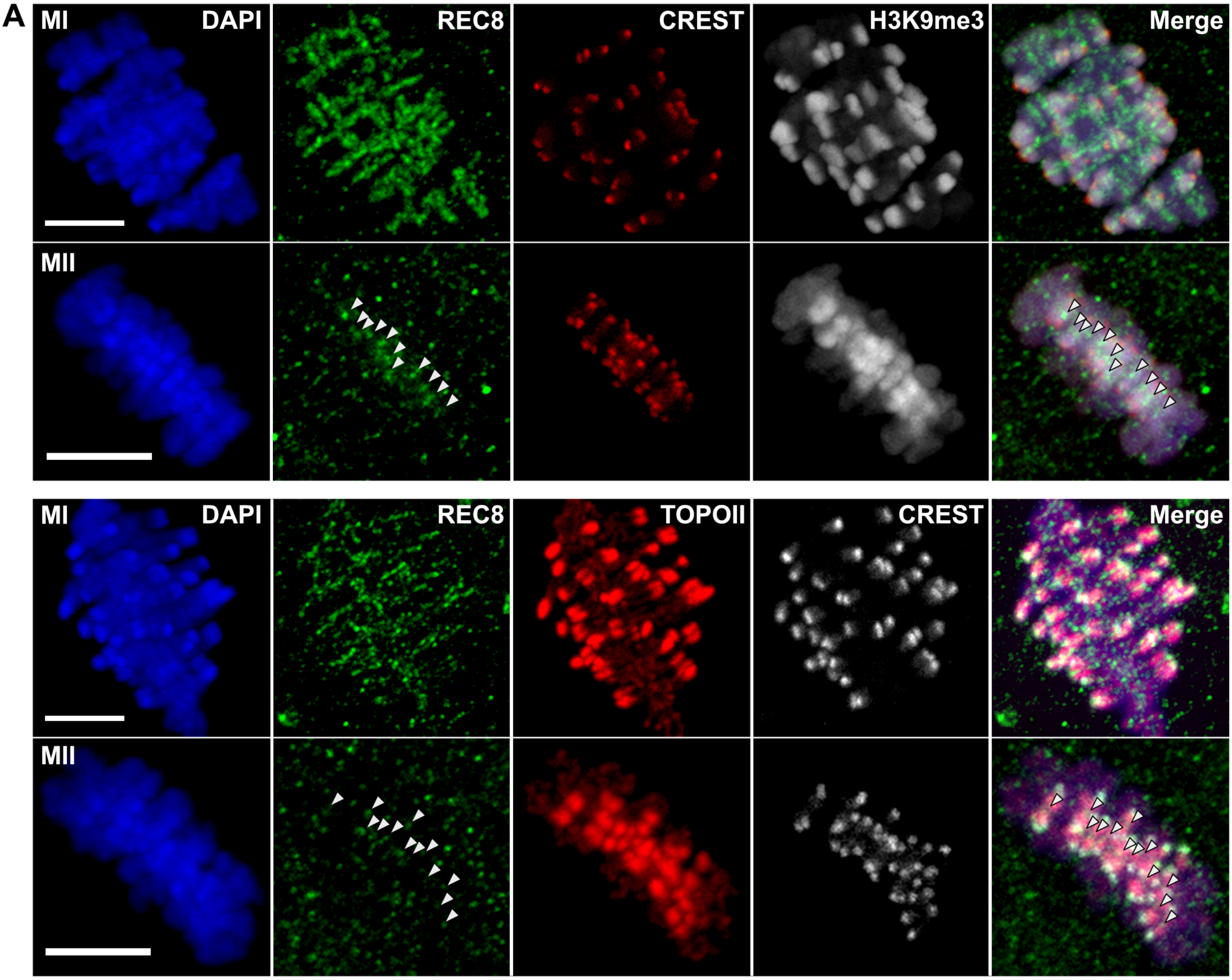

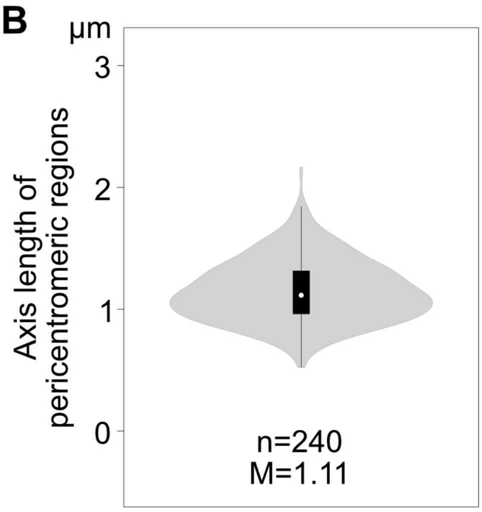

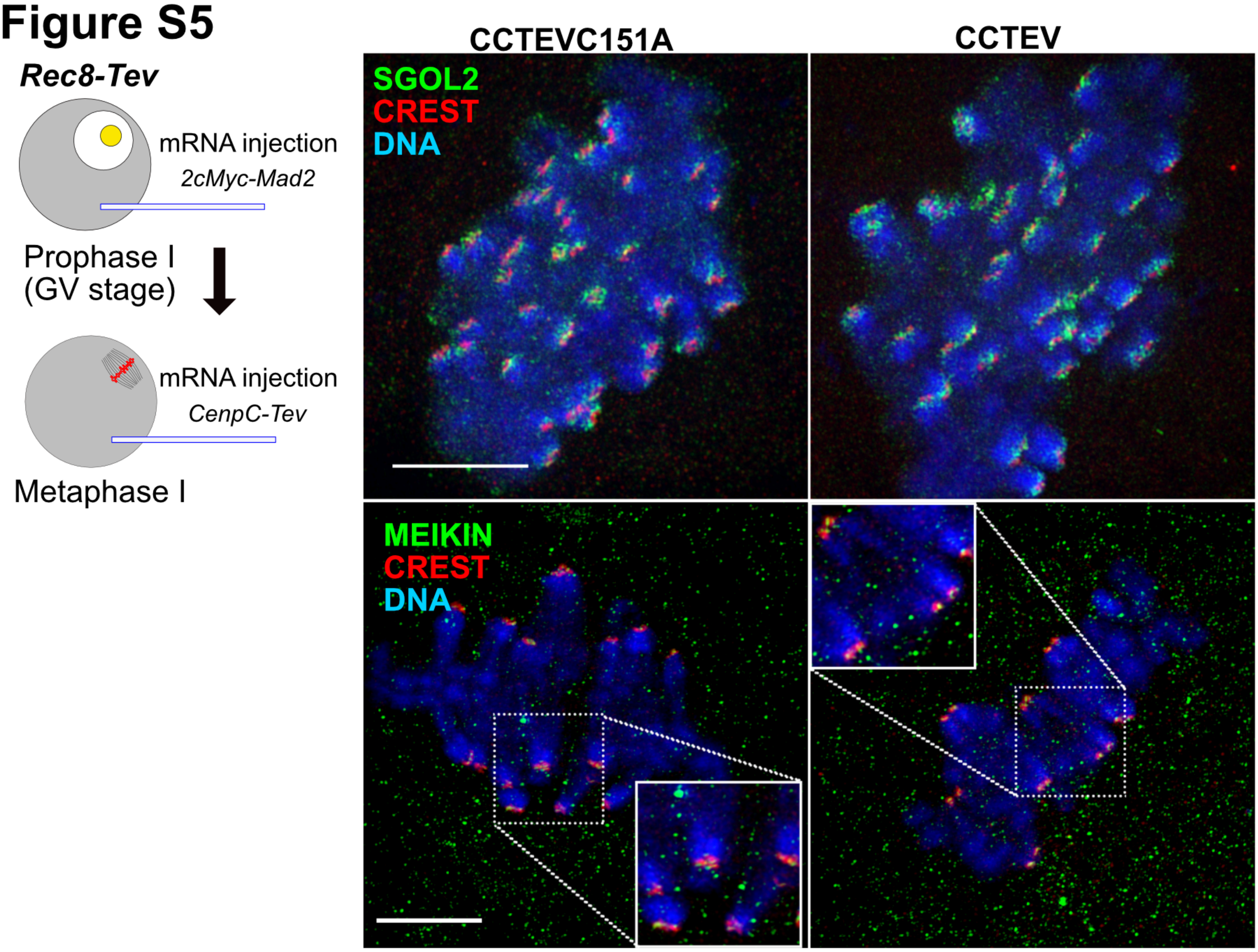

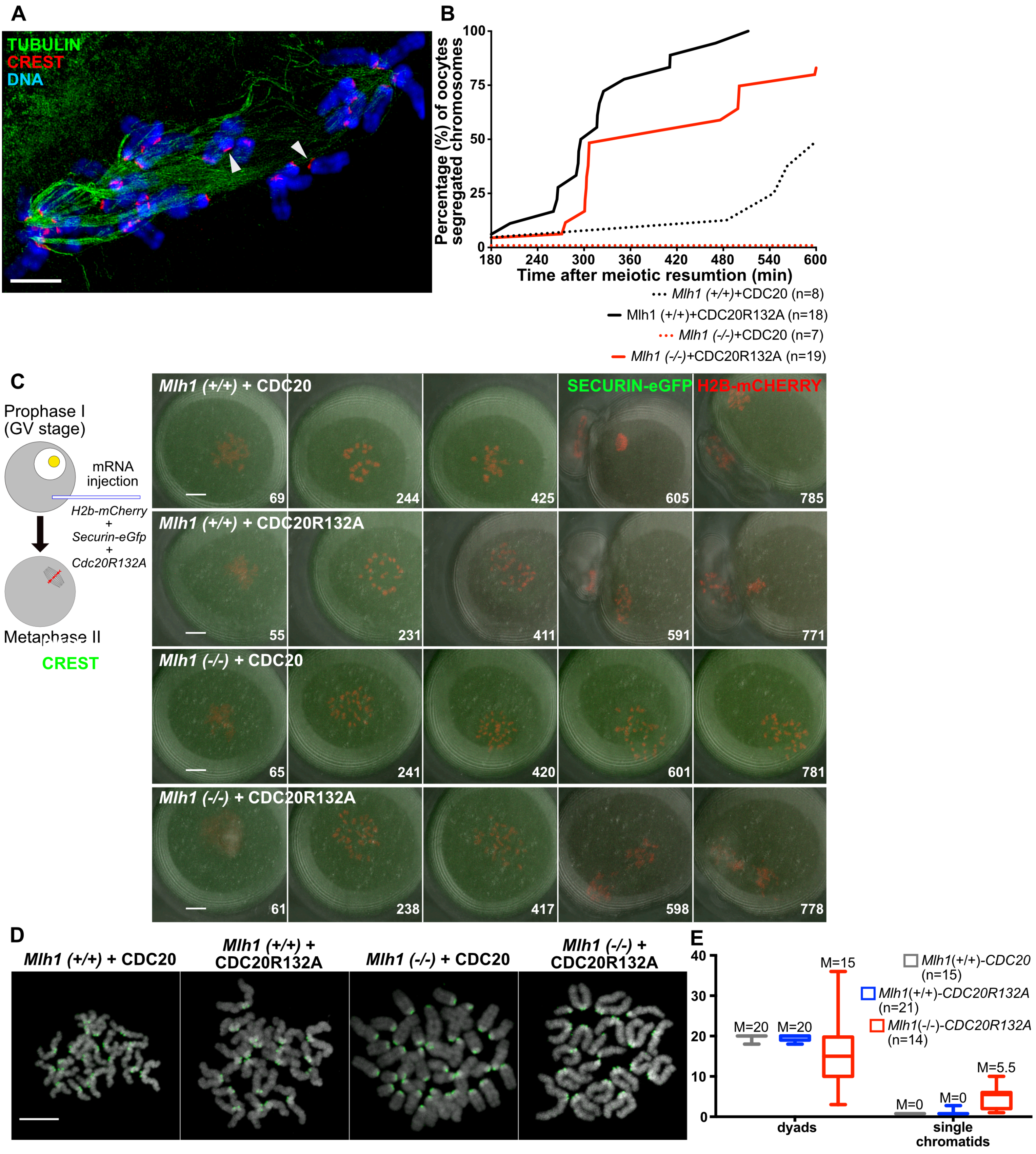

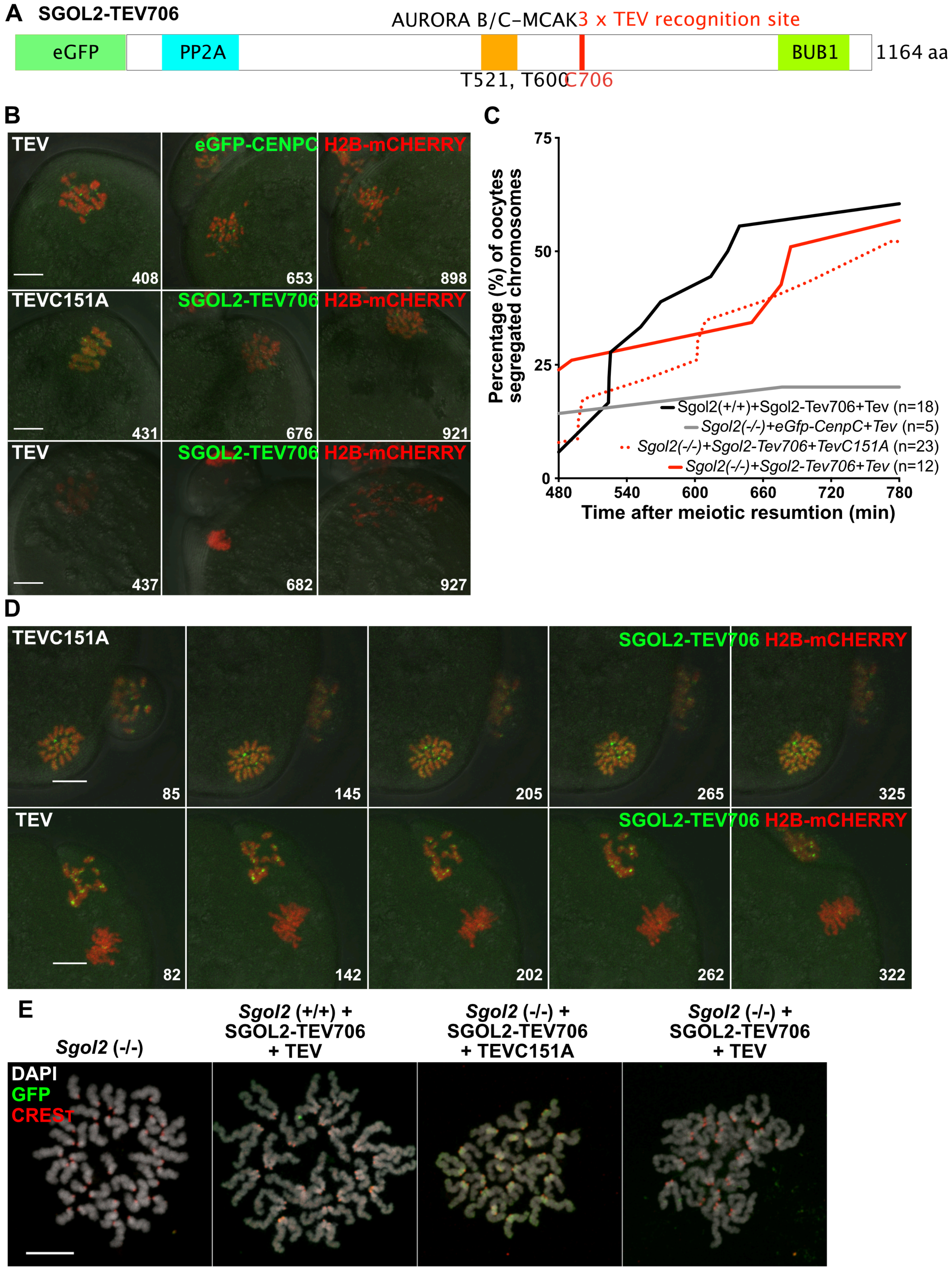
